# Genomic analysis of *Mycobacterium brumae* sustains its nonpathogenic and immunogenic phenotype

**DOI:** 10.1101/2022.12.01.518671

**Authors:** Chantal Renau-Mínguez, Paula Herrero-Abadía, Vicente Sentandreu, Paula Ruiz-Rodriguez, Eduard Torrents, Álvaro Chiner-Oms, Manuela Torres-Puente, Iñaki Comas, Esther Julián, Mireia Coscolla

## Abstract

*M. brumae* is a rapid-growing, non-pathogenic Mycobacterium species, originally isolated from environmental and human samples in Barcelona, Spain. *M. brumae* is not pathogenic and its in vitro phenotype and immunogenic properties have been well characterized. However, the knowledge of its underlying genetic composition is still incomplete. In this study, we first describe the 4 Mb genome of the *M. brumae* type strain ATCC 51384T assembling PacBio reads, and second, we assess the low intraspecies variability by comparing the type strain with Illumina reads from three additional strains. *M. brumae* genome is composed of a circular chromosome with a high GC content of 69.2 % and containing 3,791 CDSs, 97 pseudogenes, one prophage and no CRISPR loci. *M. brumae* has shown no pathogenic potential in in vivo experiments, and our genomic analysis confirms its phylogenetic position with other non-pathogenic and rapid growing mycobacteria. Accordingly, we determined the absence of virulence related genes, such as ESX-1 locus and most PE/PPE genes, among others. Although immunogenic potential of *M. brumae* was proved to be as high as *Mycobacterium bovis* BCG, the only mycobacteria licensed to treat cancer, the genomic content of *M. tuberculosis* T cell and B cell antigens in *M. brumae* is considerably lower than those of *M. bovis* BCG. Overall, this work provides relevant genomic data on one of the species of the mycobacterial genus with high therapeutic potential.

## Introduction

*Mycobacterium brumae* (synonym *Mycolicibacterium brumae*) (Meehan et al., 2021) is a saprophytic bacterium originally isolated from environmental samples, soil and water, and from human sputum from asymptomatic individuals (Luquin et al., 1993). It is a rapid-growing Mycobacterium (RGM) obtaining colonies in approximately 5-7 days at 37°C when it is grown in a rich medium. In Middlebrook 7H10 medium, *M. brumae* forms irregular, rough and non-pigmented colonies, while in Middlebrook 7H9 liquid medium can form pellicles and clumps with cording morphology (Brambilla et al., 2012).

*M. brumae* has recently been described as a plausible immunotherapy agent for non-muscle invasive bladder cancer (NMIBC) (Noguera-Ortega et al., 2016b). Currently, the standard treatment for avoiding recurrence and progression of NMIBC after transurethral resection of tumors, consists of weekly intravesical instillations of viable *Mycobacterium bovis* BCG. Despite being efficacious, BCG has some limitations. Around thirty percent of patients do not respond to BCG treatment, and a high percentage of BCG-treated patients show local and even serious systemic side effects such as pulmonary infections or sepsis (Guallar-Garrido and Julián, 2020) making adherence and continuity to treatment complicated. For that reason, safer alternatives are needed. In regards, *M. brumae* showed a safe profile in *in vitro* and *in vivo* studies (summarized in Figure 1) (Noguera-Ortega et al., 2016a; Bach-Griera et al., 2020)

**Figure 1.**
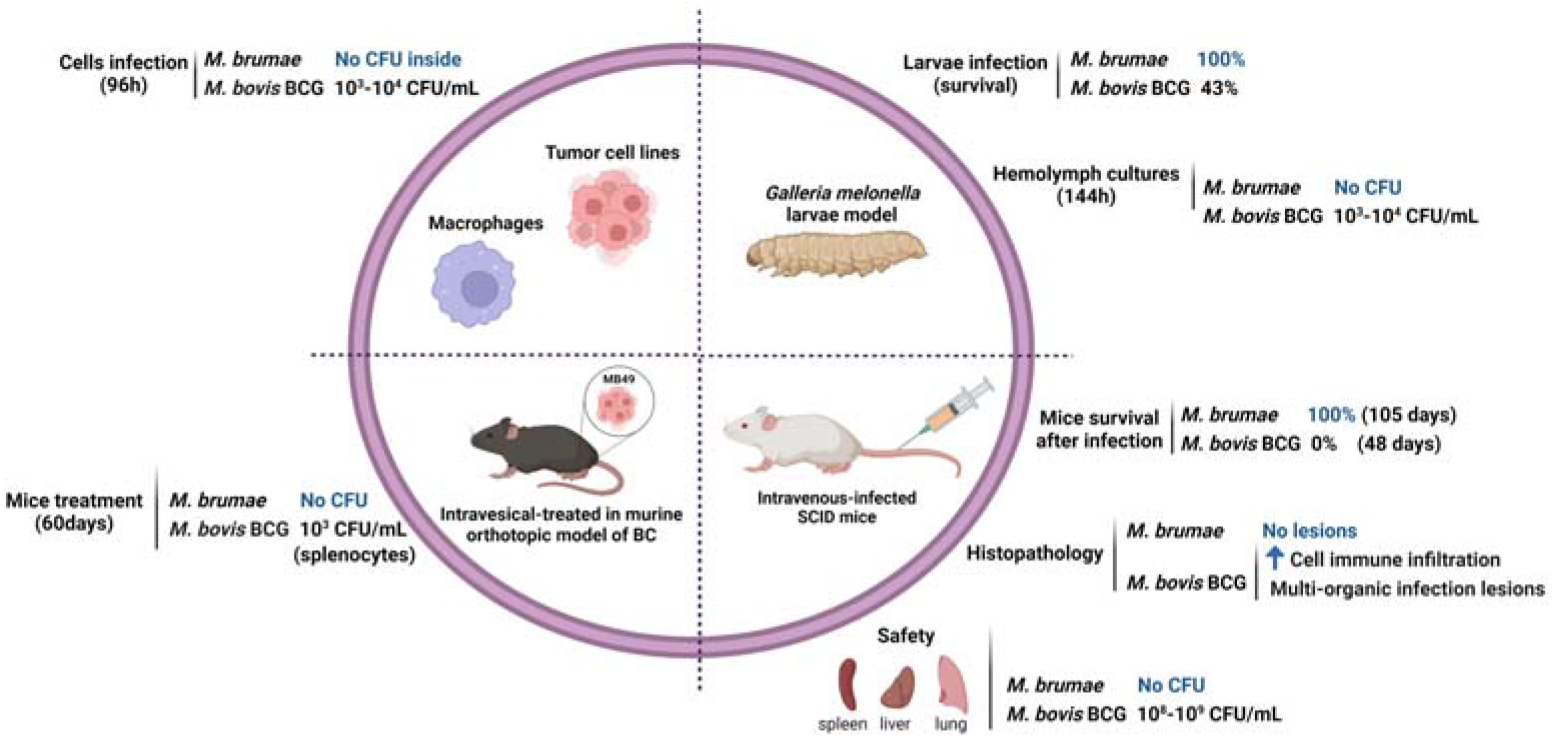
Safety profile of *M. brumae*. Comparison of the pathogenicity and toxicity between *M. brumae* and *M. bovis* BCG Connaught. Summary of the results obtained in different in vitro studies infecting macrophages (Noguera-Ortega et al., 2016b) and bladder cancer cell lines (Noguera-Ortega et al., 2016a), and in vivo studies in different animal models: intrahemacoelical infection of *Galleria mellonella* (Bach-Griera et al., 2020), intravesical instillations in orthotopic murine model of bladder cancer (Noguera-Ortega et al., 2016a, 2016b, 2016c 10.1016/j.juro.2015.07.011) and intravenous infection in SCID mice (Bach-Griera et al., 2020). Blue rising arrow (high cell immune infiltration).

Although the antitumor mechanism of action of BCG has not been fully elucidated, preclinical and clinical data supports sequential events that occur at bladder epithelium after BCG instillation. First, inhibition of tumor proliferation due to apoptosis and/or cell cycle arrest of the remaining tumor cells occurs after their interaction with BCG. In addition, BCG triggers the production of a broad spectrum of cytokines and chemokines that leads to tumor infiltration of different immune populations that indirectly provide an antitumor role. Preclinical studies have shown that *M. brumae* inhibits bladder tumor cell proliferation being even more efficacious than BCG in low-grade cells, and activated peripheral blood mononuclear cells triggering the production of cytokines and a cytotoxic profile against tumor cells (Noguera-Ortega et al., 2016a). In the orthotopic murine model of bladder cancer, the induction of local and systemic immune responses by *M. brumae* treatment leads to prolong the survival of tumor-bearing mice with respect to non-treated tumor-bearing mice and by a similar ratio than BCG treated mice (Noguera-Ortega et al., 2016a, 2016b; Guallar-Garrido et al., 2022).

However, although biological, immunological and virulence differences have been shown between *M. brumae* and BCG, it is still unknown which genomic differences might be responsible for the different biological properties of these two mycobacteria. The antigen/s responsible for the efficacious antitumor effect of BCG remain elusive. In view of the similar ability of *M. brumae* and BCG to trigger an immunomodulatory and antitumor response, it can be hypothesized that both mycobacteria share immunostimulatory antigens. Thus, it is relevant to analyze the *M. brumae* genome since it is devoid of genes involved in pathogenicity but contains genes responsible for the immunomodulatory and antitumor mechanisms. The comparison of their genomes could provide clues for understanding their therapeutic capacities. In this work we offer a detailed description of the complete genomic sequence of the reference strain of *M. brumae*, and we address the biological differences between *M. brumae* and BCG in the context of their genomic differences using a comparative genomics approach. Furthermore, we assess the intraspecies variability of *M. brumae* by comparing the genetic diversity between different isolates of the species.

## Materials and Methods

### Bacterial strains

Four *M. brumae* strains (CR103, CR142, CR269, and CR270 (which corresponds to the type strain: ATCC 51384T)) were provided by Prof. Luquin, obtained from the original isolates described in Luquin et al. (1993). Lyophilized cells were cultured on Middlebrook 7H10 agar medium (Difco Laboratories, Michigan, USA) supplemented with 10% oleic-albumin-dextrose-catalase (OADC) and incubated at 37°C for one week.

### Genomes

We analyzed the phylogenetic relationships of published reference/complete genomes from different *Mycobacterium* species (Supplementary Table S1). Although the genome sequence is not closed, three additional previously published *M. brumae* sequences for the type strain were compared with our reference genome sequence: CIP1034565 (GCF_002553575.1), DSM44177 (GCF_004014795.1) and MBR1 (GCF_900073015.1).

The *Mycobacterium tuberculosis* H37Rv and *M. bovis* BCG Connaught genomes used for genome comparisons were downloaded from the GenBank database with accession numbers GCA_000195955.2 and GCA_001287325.1 respectively.

### Drug resistance analysis

Sensititre RAPMYCOI and SLOMYCOI panels (Thermo Fisher Scientific, Massachusetts, USA) were used to determine the susceptibilities of *M. brumae* strains to different antibiotics. A wide variety of antibiotic groups were tested, including β-lactams (amoxicillin/clavulanic acid, cefepime, cefoxitin, ceftriaxone, imipenem), aminoglycosides (amikacin, streptomycin, tobramycin), quinolones (ciprofloxacin, moxifloxacin), macrolides (clarithromycin), tetracyclines (doxycycline, minocycline, tigecycline), oxazolidinones (linezolid), sulfonamides (trimethoprim/sulfamethoxazole) and various anti-tuberculosis drugs (ethambutol, ethionamide, isoniazid, rifabutin, rifampin). A bacterial suspension adjusted to a McFarland 1 standard was prepared, and 50 μL of the suspension was transferred to a tube of Mueller-Hinton broth, obtaining an inoculum of 5×105 CFU/mL. 100μL of bacterial inoculum was added to each well of the Sensititre plates. The rest of the procedure was done according to the manufacturer’s instructions. The method and guidelines for the interpretation of results were those of the Clinical and Laboratory Standards Institute. The antimicrobials p-aminosalicylic acid, capreomycin, cycloserine and kanamycin susceptibility results were obtained from previously published studies (Luquin et al., 1993).

The reported mutations in *M. tuberculosis* H37Rv associated with drug resistance were obtained from the WHO 2021 annual catalog of mutations in *M. tuberculosis* (Catalogue of mutations in *Mycobacterium tuberculosis* complex and their association with drug resistance, 2021) accessed in April 2022). The presence or absence of the mutations in the regions of interest *(rpoB* and *katG* genes and *inhA* regulation regions) were manually inspected in the gene sequence extracted from in the *M. brumae* genome using MEGA X: Molecular Evolutionary Genetics Analysis across computing platforms (Tamura et al., 2021).

### DNA extraction

DNA was extracted from the four *M. brumae* strains using the UltraClean® Microbial DNA Isolation Kit following the manufacturer’s instructions (MO BIO Laboratories, Inc. Carlsbad, California, USA) with slight modifications. Briefly, samples were heated to 65°C for 10 minutes in step 4 of the procedure, to improve production. The extracts obtained were electrophoresed on a 0.8% agarose gel using 6x NZYDNA Loading Dye (Nzytech, Lisboa, Portugal) and λ DNA/Hind III marker (Thermofisher, Massachusetts, USA). Samples were concentrated by evaporation at 45°C with Eppendorf® centrifugal vacuum concentrator 5301 (Sigma-Aldrich, Missouri, USA) and DNA quantification was performed using a NanoDropTM 2000 spectrophotometer (NanoDrop Technologies, Inc. Wilmington, USA).

### Library construction and genome sequencing

Whole-genome sequencing for the *M. brumae* ATCC 51384T type strain was performed at the SCSIE Genomics Core Facility at the University of Valencia using PacBio Sequel™ system (Pacific Biosciences, Menlo Park, CA, USA). *M. brumae* gDNA concentration was measured by Qubit dsDNA HS kit (ThermoFisher Scientific, Mississauga, ON, Canada). The library was prepared following the manufacturer’s protocol for 10□kb SMRTbell Express Template Prep Kit 2.0 (Pacific Biosciences). Briefly, genomic DNA (2□μg) was diluted in 200 μL of elution buffer and fragmented using g-TUBE (Covaris) by centrifuging at 2400×g for 2□min. The fragmented DNA was concentrated using AMpure PB magnetic beads (Beckman Coulter) and used for the preparation of a single non-multiplexed sequencing library. A ready-to-sequence SMRT bell-Polymerase Complex was created using PacBio’s Sequel binding kit 3.0 according to the manufacturer’s instructions. The final library was sequenced on 1 Sequel™ SMRT® Cell 1□M v3, taking a 10□h movie using the Sequel Sequencing Kit 3.0. *M. brumae* strains CR103, CR142 and CR269 were subjected to whole-genome sequencing with Illumina. Genome libraries were constructed with a Nextera XT DNA library preparation kit (Illumina, San Diego, CA), following the manufacturer’s instructions. Sequencing was carried out at Institute of Biomedicine of Valencia Ion an Illumina MiSeq platform (2×300 cycles paired-end reads).

### Assembly and annotation of the completed genome

Sequence assembly was performed from 584,529,485 total bases (sequencing raw data used for the assembly is available in NCBI under the code PRJNA798885) with a read depth of 136 over the genome and read length N50 of 6,188 bases using SMRT Link v8.0.0 interface and Microbial Assembly analysis application (Pacific Biosciences). Genome assembly was conducted with genome size parameter set to 4 Mb and produced a single polished circular contig of 3,988,920 bases, and the quality of the assembly was evaluated using Inspector software (Chen et al., 2021), by mapping the long reads to the contig. Ori-Finder 2022 (https://tubic.org/Ori-Finder2022, accessed in August 2022) was used to identify the bacterial replication origin. Draft genome for the *M. brumae* ATCC 51384T strain was annotated by the pipelines Rapid Prokaryotic Genome Annotation (Prokka) v 1.14.5 (Seemann, 2014), PGAP v 2022-04-14.build6021 (Tatusova et al., 2016), Bakta v1.4.2 (Cantalapiedra et al., 2021) and the Rapid Annotations using Subsystems Technology (RAST) server (Aziz et al., 2008). PseudoFinder (Syberg-Olsen et al., 2021) and PGAP were used for the detection of pseudogene candidates. Only those pseudogenes predicted by both approaches were kept as highly confident pseudogenes, while the rest were manually annotated as “putative pseudogene”. PGAP annotation was curated by detailed inspection of virulence related genes detailed in next section. Additionally, all ESX regions were obtained from *M. tuberculosis* (Mycobrowser, accessed in April 2022) and BLASTp was used to identify and manually annotate those regions in our *M. brumae* genome. For the identification of other regions of interest in our genome, CRISPRFinder (Grissa et al., 2007) was used for the CRISPR-Cas system prediction. The internal conservation of the candidate DRs and the divergence of the candidate spacers were manually checked. PhiSpy v 4.2.21 was used to identify the prophages harbored in the *M. brumae* genome. CIRCOS software (Krzywinski et al., 2009) was used to represent the circular representation of the chromosome. To compare the functional categories between *M. brumae* and *M. tuberculosis* H37Rv, the COG assignment for each *M. brumae* protein was performed by EggNOG-mapper v 2.1.9 (Cantalapiedra et al., 2021) and *M. tuberculosis* COG assignments were obtained from the COG database (https://www.ncbi.nlm.nih.gov/research/cog). The abundance of proteins in each category was compared. Row-wise Fisher’s exact test was used within R v 4.2.1 to assess the statistical significance of differences between the *M. brumae* and *M. tuberculosis* H37Rv annotations for the gene abundance in each COG category.

### Virulence genes analysis

To obtain a list of virulence genes, we looked for virulence-related genes from *M. tuberculosis* H37Rv in *M. brumae* and in *M. bovis* BCG Connaught. The list of genes was compiled by combining 249 genes included in the Virulence Factors Database (VFDB) (http://www.mgc.ac.cn/VFs/, accessed in July 2022) and 268 additional genes reported to be involved in virulence in vivo or in vitro by reviewing the bibliography. Because 193 genes were retrieved by both approaches, a final list of 324 genes was analyzed. The bibliographic search was done looking for in PubMed the following terms “((virulence[Title/Abstract]) AND (Mycobacterium tuberculosis[Title/Abstract])) AND ((in vivo[Title/Abstract]) OR (in vitro[Title/Abstract]))”. We retrieved 594 manuscripts and included those which studied a virulence phenotype with *M. tuberculosis*, obtaining a list of 199 articles. The complete list of references consulted for the virulence-related genes is indicated in Supplementary Table S2.

To assess the presence or absence of these *M. tuberculosis* genes in other genomes, we first retrieved the nucleotide sequences from H37Rv genomes (accession number GCA_000195955.2) by indexing the nucleotide position with Python module Biopython version 1.79 (Cock et al., 2009), and retrieved the coordinates and orientation of the genes from Mycobrowser (https://mycobrowser.epfl.ch/). Next, we translated and obtained the amino acid sequence. For genes in the negative strand, we performed the complementary and reverse translation by using the Biopython module. tBLASTn was then used to find the amino acid similarity between *M. tuberculosis* proteins and both *M. bovis* BCG Connaught and *M. brumae* proteins using BLAST+ v 2.10.0 (Camacho et al., 2009) by a custom Python script v 3.6.0 (GitHub repository: https://github.com/PathoGenOmics/mbrumae_closedgenome). After exploring different percentages of identity, a gene was considered present when 70% identity and 70% coverage were reached. Additionally, similarity between *Mmpl, Mce* and PE/PPE proteins from our *M. brumae* genome (CP104302) and previously published M. brumae contigs (GCF_002553575.1) and *M. fallax* (GCA_010726955.1) was assessed using the same procedure explained above.

### Immunogenic genes analysis

For the analysis of immunogenic genes, all the experimentally verified T and B cell epitopes were obtained from the Immune Epitope Database for *M. tuberculosis* H37Rv (IEDB.org: Free epitope database and prediction resource, accessed in April 2020) studied in humans in the context of infectious diseases, allergy, autoimmunity, and transplantation (Supplementary Table S2). Then, the immunogenic epitope-associated genes were obtained from *M. tuberculosis* H37Rv using a custom pipeline (https://github.com/PathoGenOmics/mbrumae_closedgenome). The procedure followed to assess the presence of a gene was the same as the explained above for the virulence-related genes.

### Cell wall biosynthesis genes analysis

To obtain a list of *M. tuberculosis* genes involved in the cell wall biosynthesis the following bibliographic search was done in PubMed: “(Mycobacterium tuberculosis[Title/Abstract]) AND (biosynthesis[Title/Abstract]) AND ((mycolic acids[Title/Abstract]) OR (PDIM[Title/Abstract]) OR (PGL[Title/Abstract]) OR (PIM[Title/Abstract]) OR (TDM[Title/Abstract]) OR (TMM[Title/Abstract]))”. We obtained a list of 221 articles, and we filtered for studies that described genes with cell-wall related function in *M. tuberculosis*, obtaining a final list of 95 articles and 179 genes. The complete list of references consulted for the cell wall genes is indicated in Supplementary Table S2. The procedure followed to assess the presence of a gene was the same as the explained above for the virulence-related genes.

### Mapping and variant calling in Illumina sequences

Two different pipelines for mapping and variant calling were used for the analysis of intraspecific variability of three different *M. brumae* strains respect to the reference genome. We used two pipelines previously described (Chiner-Oms et al., 2019) (Coscolla et al., 2021). Briefly, the FASTQ were trimmed with Trimmomatic v 0.33 (SLIDINGWINDOW 5:20) to remove Illumina adaptors and low-quality reads (Bolger et al., 2014), excluding from downstream analysis the ones shorter than 20 bp. Overlapping paired-end reads of 15 nucleotides size were merged with SeqPrep v1.2 (https://github.com/jstjohn/SeqPrep, accessed in 2020). The resultant reads from the three *M. brumae* strains (CR103, CR142, and CR269) were aligned to the *M. brumae* ATCC 51384T complete genome obtained using BWA v 0.7.13 (mem algorithm) (Li and Durbin, 2010). Duplicated reads were marked by the Mark Duplicates module of Picard v 2.9.1 (https://github.com/broadinstitute/picard, accessed in 2020) and excluded. Reads with an alignment score corresponding to more than 7 mismatches per 100 bp were excluded using Pysam v 0.9.0 to avoid false positives (https://github.com/pysam-developers/pysam, accessed in 2020). SNPs were called with SAMtools v 1.2 mpileup (Li, 2011) and VarScan v 2.4.1 (Koboldt et al., 2012). SNPs were considered only if the variant allele frequency was higher than 10%, i.e., variants that were detected in at least 10% of the reads for that position. Only SNPs that reached fixation within an isolate were considered as homozygous SNPs (within-host frequency, i.e., SNP frequency within the reads of the same sample, higher 90%), being classified as heterozygous SNPs otherwise. All SNPs were annotated using SnpEff v 4.11 (Cingolani et al., 2012), in accordance with the *M. brumae* ATCC 51384T strain annotation (CP104302). Repeated regions in the *M. brumae* genome were obtained using run-mummer3 and a custom pipeline using Python to identify repeated genes with a minimum threshold of 300 bp of identity. The custom script used and the list of repeated genes were uploaded to Github (https://github.com/PathoGenOmics/mbrumae_closedgenome). SNPs located in those regions were classified as “low confidence SNPs”. For the deletions analysis, mean coverages per gene corrected by the size of the gene were calculated (https://github.com/PathoGenOmics/mbrumae_closedgenome). Only those genes with at least 25% of the gene length with less than 5 reads-per-site were considered as deleted. As repetitive regions could compromise the accuracy of the analysis, genes located in repeated regions were also excluded.

### Phylogenetic reconstruction

For the main phylogenetic construction, we selected a manageable subset of genomes from the Mycobacterium genus that would represent different parts of the genus, and specially the closest taxa to *M. brumae*. We only used closed and well-described genomes (Fedrizzi et al., 2017). Our dataset included 21 downloaded genome assemblies (Supplementary Table S1), our newly sequenced *M. brumae* assembly and the CR103, CR142 and CR269 *M. brumae* strains. Roary software v 3.11.2 (Page et al., 2015) was used to obtain the aligned core-genome, defined as the subset of genes found in at least 99% of the samples, and included 177 conserved genes (Supplementary Table S3). The alignment was used to infer a maximum-likelihood phylogenetic tree using IQ-TREE v 1.6.12 (Nguyen et al., 2015) with GTR model and the tree was re-examined using the bootstrap (1000 replicates) resampling method. Hoyosella subflava (family Mycobacteriaceae) was used to root the tree as it is the sequenced bacterium closest to the Mycobacterium genus (Gupta et al., 2018). Virulence potential and growth rate (Fedrizzi et al., 2017) was annotated on the phylogeny using the iTOL software v 4 (Letunic and Bork, 2019). Possible recombination events in the tree were checked using Gubbins v 3.2 (Croucher et al., 2015).

An additional phylogenetic analysis was performed to confirm the consistency of our genomic sequence of *M. brumae* reference genome with other genomic sequences of the same strain. We included *M. brumae* ATCC 51384 sequence, the three Illumina strains (CR103, CR142 and CR269) and three published *M. brumae* sequences for the type strain which are not closed (named in Supplementary Figure S1 as CIP1034565, DSM44177 and MBR1 with accession numbers GCF_002553575.1, GCF_004014795.1 and GCF_900073015.1 respectively). Panaroo software v 1.2.9 (Tonkin-Hill et al., 2020) was used to obtain the core-genome, using the following parameters “panaroo -i *.gff -o results_panaroo_core --clean-mode strict -a core”, and the alignment was constructed using IQ-TREE software v 1.6.10 with GTR model and the tree was re-examinated using the bootstrap (1000 replicates) resampling method.

## Results

### Genome composition of the reference strain

To gain further insights into its genome composition and intraspecific variation, the complete *M. brumae* reference genome sequence was determined by PacBio. The complete genome from *M. brumae* and the manually curated annotation has been deposited in the GenBank database under the BioProject code PRJNA798885 with accession number CP104302. The *M. brumae* genome consisted of a single circular chromosome of 3,988,920 bp with an average GC content of 69% (Table 1). The quality of the assembly was assessed by a quality value score of 63 using Inspector software (Chen et al., 2021). The oriC was predicted next to the rnpA-rpmH-dnaA-dnaN-recF-gyrB-gyrA region (Supplementary Table S4) and showed in Figure 2. Conversely, the GC skew did not accurately indicate the oriC (Figure 2). Different annotations of *M. brumae* resulted in an estimate of 3,791, 3,794, 3,827 and 3,781 protein-coding sequences (CDS) using PGAP, Bakta, RAST and Prokka respectively (Supplementary table S4). Because PGAP delivered fewer CDS as hypothetical proteins (Table 1), we kept the PGAP annotation for manual curation (Supplementary Table S4). Additionally, we found 48 genes encoding tRNA and two rRNA operons, each one including three rRNAs.

**Figure 2.**
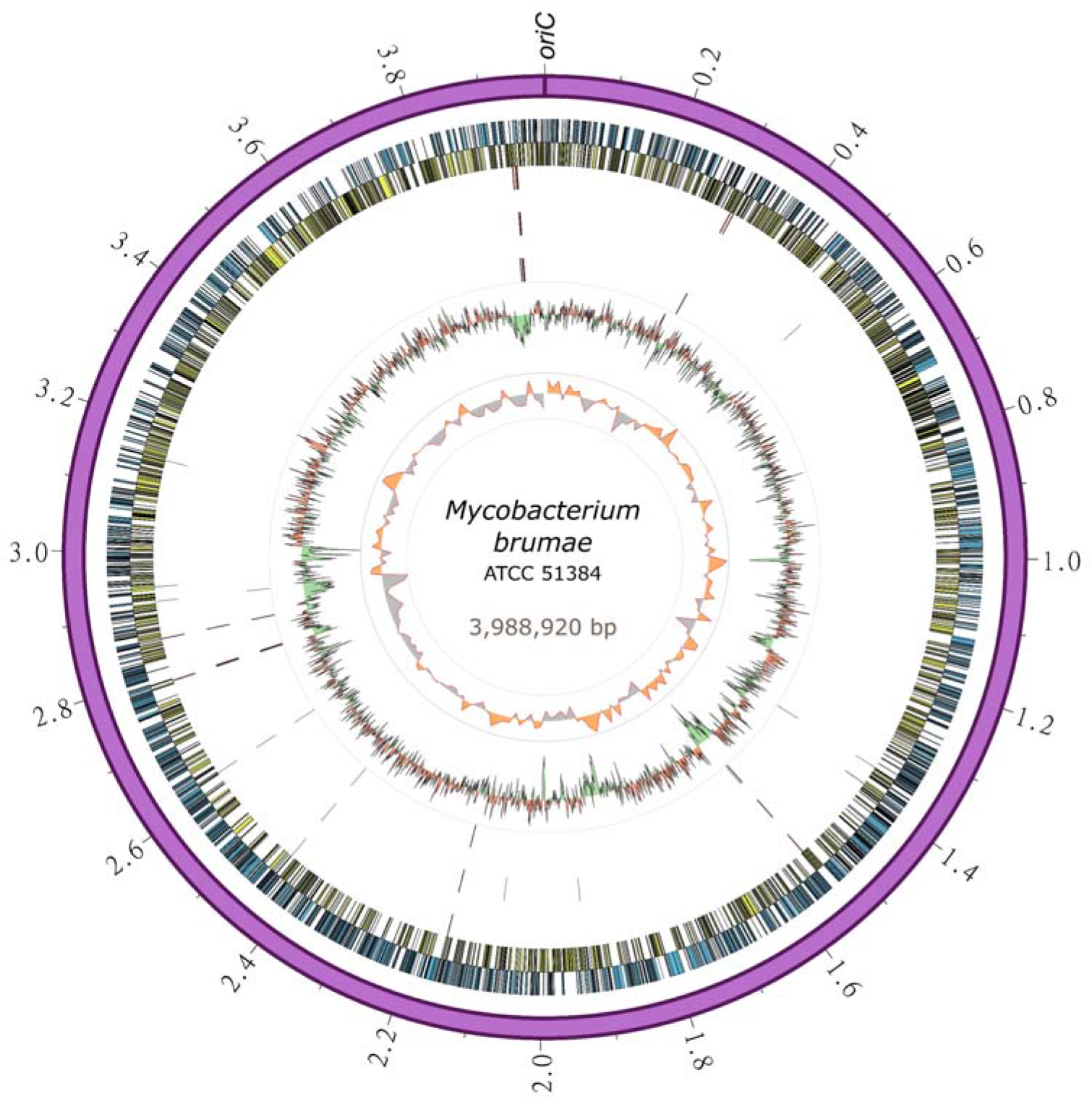
Circular representation of the *M. brumae* chromosome. The outermost ring shows the chromosome length in megabases, followed by the forward (blue) and reverse genes (yellow). SNPs in *M. brumae* strains (CR269, CR142 and CR103) with respect to the ATCC 51384T reference genome is indicated in the next three rings respectively. Moving inward, the next ring shows the GC content, with values over the mean filled in red and values below the mean in green. The last ring shows the GC skew (G-C)/(G+C) using a 20-kb window.

**Table 1.**
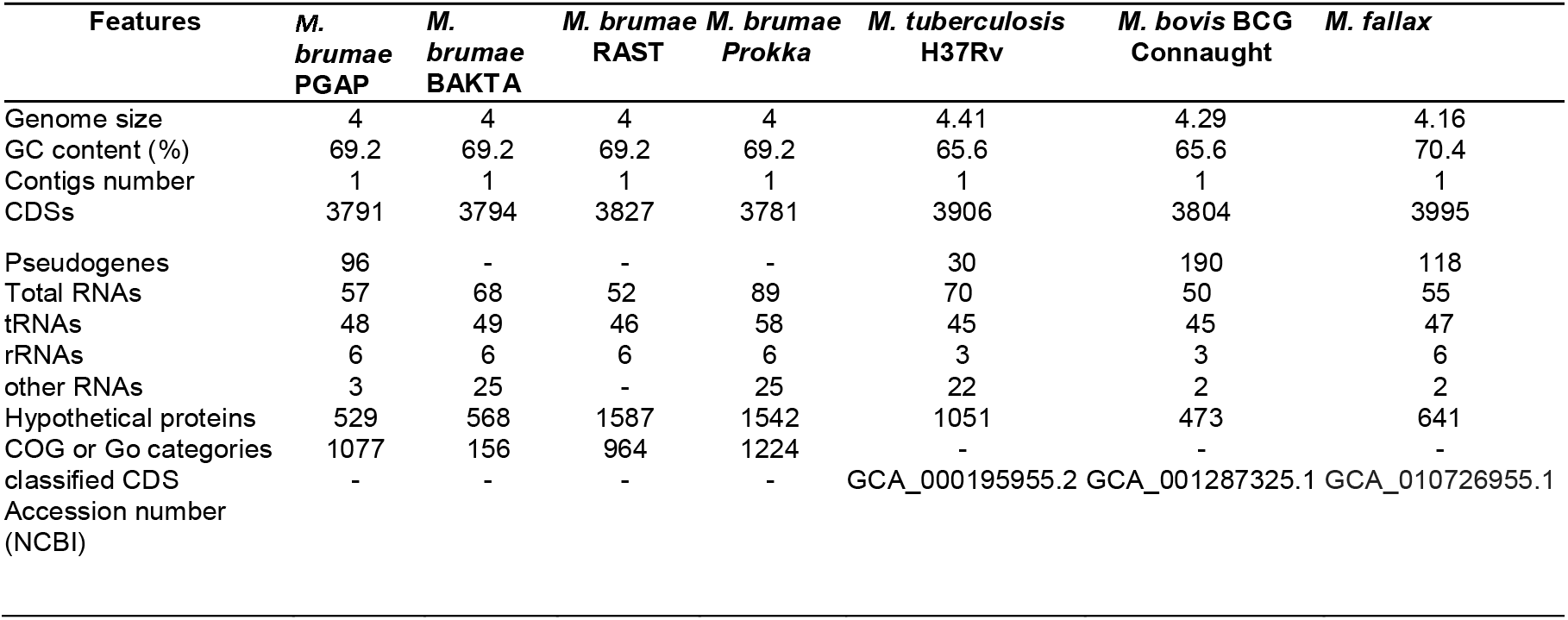
Genome features. Genome overview of *M. brumae* based on PGAP, Bakta, RAST and Prokka annotations and compared with *M. tuberculosis* H37Rv, *M. bovis* BCG Connaught and *M. fallax*.

| Features | *M. brumae*<br>PGAP | *M. brumae*<br>BAKTA | *M. brumae*<br>RAST | *M. brumae*<br>Prokka | *M. tuberculosis*<br>H37Rv | *M. bovis* BCG<br>Connaught | *M. fallax* |
| --- | --- | --- | --- | --- | --- | --- | --- |
| Genome size | 4 | 4 | 4 | 4 | 4.41 | 4.29 | 4.16 |
| GC content (%) | 69.2 | 69.2 | 69.2 | 69.2 | 65.6 | 65.6 | 70.4 |
| Contigs number | 1 | 1 | 1 | 1 | 1 | 1 | 1 |
| CDSs | 3791 | 3794 | 3827 | 3781 | 3906 | 3804 | 3995 |
| Pseudogenes | 96 | - | - | - | 30 | 190 | 118 |
| Total RNAs | 57 | 68 | 52 | 89 | 70 | 50 | 55 |
| tRNAs | 48 | 49 | 46 | 58 | 45 | 45 | 47 |
| rRNAs | 6 | 6 | 6 | 6 | 3 | 3 | 6 |
| other RNAs | 3 | 25 | - | 25 | 22 | 2 | 2 |
| Hypothetical proteins | 529 | 568 | 1587 | 1542 | 1051 | 473 | 641 |
| COG or Go categories | 1077 | 156 | 964 | 1224 | - | - | - |
| classified CDS | - | - | - | - | GCA_000195955.2 | GCA_001287325.1 | GCA_010726955.1 |
| Accession number<br>(NCBI) |  |  |  |  |  |  |  |

Pseudogene annotation was performed by combining the results from two approaches. PGAP predicted 97 and PseudoFinder predicted 326 pseudogenes, of which 47 were predicted by both. In order to be conservative, we annotated those detected by both approaches as pseudogenes and those detected just by PGAP as probable pseudogene (Supplementary Table S4). The number of pseudogenes found with both approaches was similar to the number of pseudogenes in *M. tuberculosis* (GCA_000195955.2) and lower than those found in *M. bovis* (GCA_001287325.1) or *M. fallax* (GCA_010726955.1) (Table 1). Another interesting feature in the genome refers to the bacterial adaptive systems against viral infections. We analyzed the clustered regularly interspaced short palindromic repeats (CRISPRs) and prophages. The CRISPRfinder software predicted that there were no CRISPR sequences throughout the *M. brumae* genome. Next, we used the PhiSpy software to identify the prophages present in the genome, and one insert was detected between the positions 83324-110205 within the *M. brumae* genome, including a total of 235 CDS (Supplementary Table S5).

For a general genome content description, CDS predicted by PGAP annotation were subclassified into 26 different COG categories (Figure 3) and compared to *M. tuberculosis*. We found 2617 (69%) *M. brumae* CDSs associated with a COG category, including 661 (25%) assigned to less informative categories such as “General function prediction only” and “Function unknown” (Figure 3). Compared to the *M. tuberculosis* annotation, *M. brumae* had a significantly bigger representation in L (“Replication and repair”) and M (“Cell wall/membrane/envelope biogenesis”) categories with 6.57% and 7.87% of the genes in *M. brumae* compared to 3.35% and 4.97% in *M. tuberculosis* (p-values of 4.70e-07 and 1.86e-04 respectively). *M. brumae* was also more abundant (8.06%) than *M. tuberculosis* (6.18%). On the contrary, *M. tuberculosis* showed a statistically higher abundance of genes in six categories compared to *M. brumae*, including lipid metabolism (8.75% and 4.47% for *M. tuberculosis* and *M. brumae* respectively, p-value 3.08e-09), coenzyme metabolism (6.63% and 3.59%, p-value of 2.80e-06), mobilome (2.67% and 0%, p-value of 3.13e-21), secondary structure (5.61% and 3.52%, p-value of 2.03e-04), defense mechanisms (3.28% and 1.91%, p-value of 1.42e-03) and signal transduction (3.51% and 2.18%, p-value of 2.97e-03).

**Figure 3.**
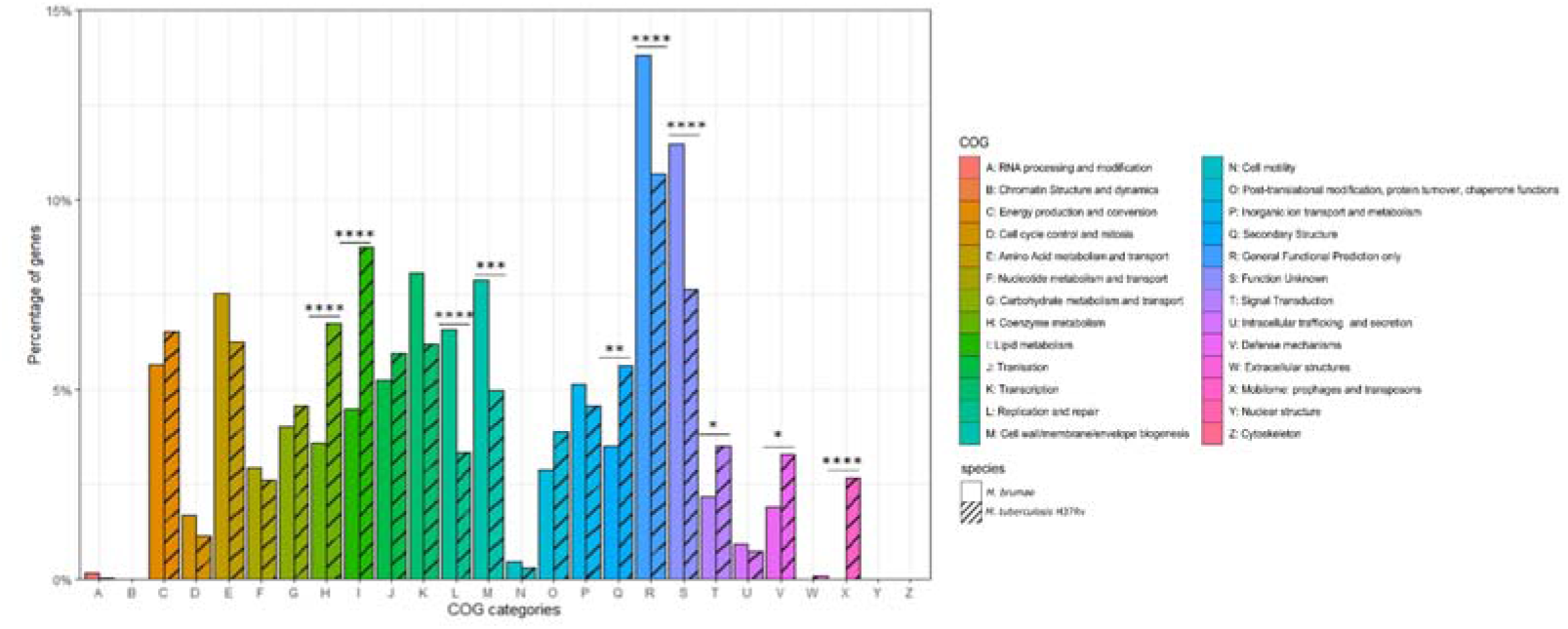
Gene annotation comparison of COG distributions shared in *M. brumae* and in *M. tuberculosis* H37Rv. COG refers to Clusters of Orthologous Groups. The vertical axis shows the percentage of genes in each category. The different categories are represented on the horizontal axis and the legend indicates the correspondence with each COG category. The corresponding strain for each bar is indicated in stripe pattern for *M. tuberculosis* H37Rv and in plain pattern for *M. brumae*. Fisher test p-values for each category indicated as following: *p<0.05; **p<0.01; ***p<0.001; ****p<0.0001.

### Phylogenetic position of *M. brumae* in the genus

To investigate the *M. brumae* position in the evolutionary history of mycobacteria, we constructed a phylogeny using a representative sample of species from the genus (Figure 4). The alignment was constructed using all the core genes among the different species to provide a higher resolution of the relationships among them. Recombination events were eliminated from the alignment and the structure of the tree remained unaltered, which supports the robustness of the analysis. *M. brumae* clusters with other non-pathogenic rapidly growing mycobacteria. *M. fallax* was the closest mycobacteria to *M. brumae*, followed by *Mycobacterium insubricum*, and all three clustering with the monophyletic group formed by *Mycobacterium confluentis* and *Mycobacterium chitae*.

**Figure 4.**
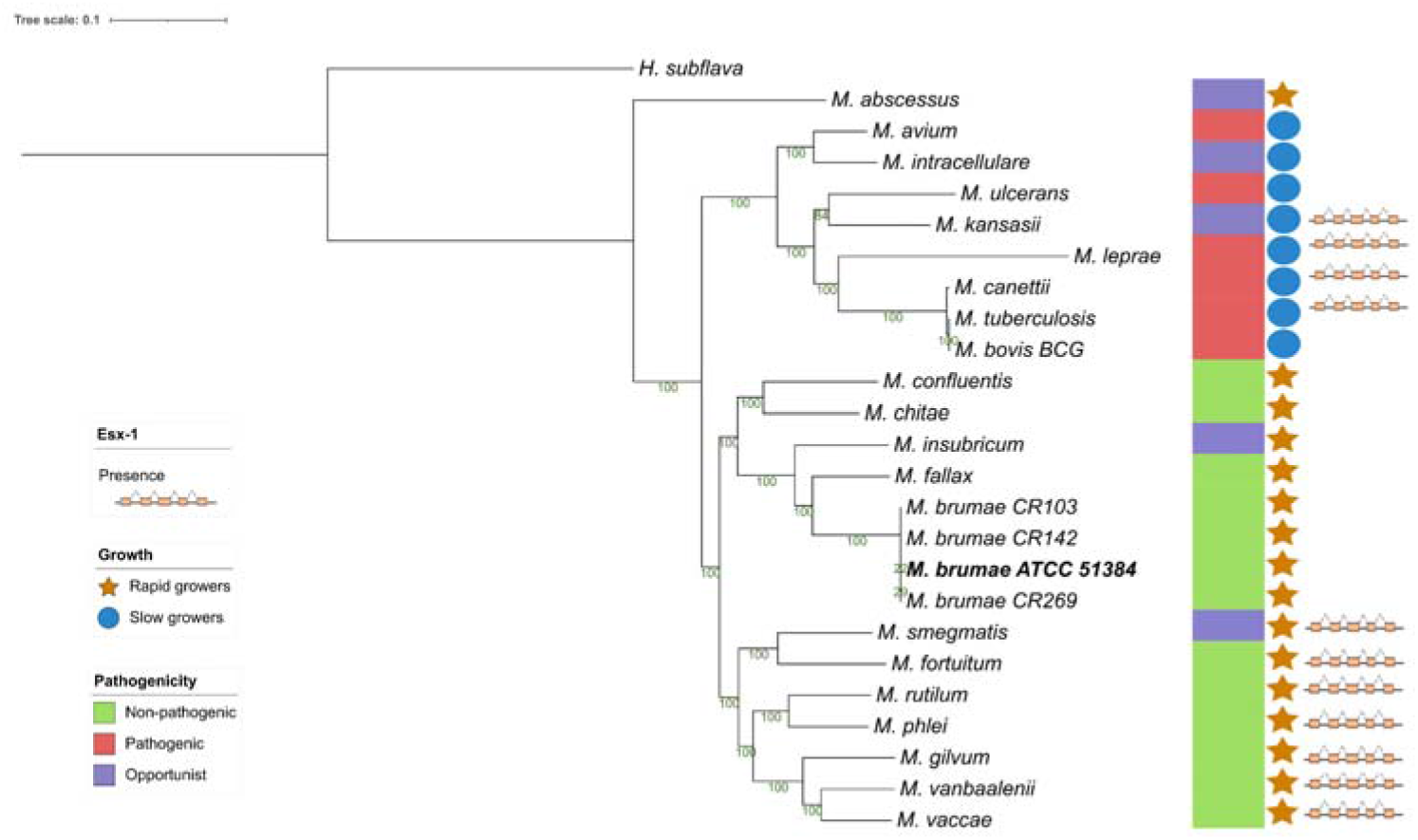
Phylogenetic relationship of *M. brumae* within the *Mycobacterium* genus. Maximum likelihood phylogenetic tree of type strains from the *Mycobacterium* genus based on an alignment with 177 core genes, highlighting the position of *M. brumae* ATCC 51384^T^ type strain. GTR model was used with bootstrap confidence of 1000 replicates. Branch lengths are proportional to nucleotide substitutions and the topology is rooted with *H. subflava*. The phylogenetic tree was annotated with the pathogenicity, the ESX-1 presence or absence, and the growth rate (see Methods).

Although the genome presented here is the first *M. brumae* closed genome, the same strain has been sequenced previously. We assessed the consistency of our closed genome sequence with previously We assessed the consistency of our closed genome sequence with previously published *M. brumae* type strain sequences. Phylogenetic analysis shows that our *M. brumae* sequence clustered with the monophyletic group formed by MBR1 and CIP1034565, all of them corresponding to the type strain (Supplementary Figure S1). Another monophyletic group was formed by the other three strains (CR103, CR142 and CR269) and DSM44177 which also corresponds to the type strain. This result indicates the consistency of our results with two out of three genomic sequences of the same strain.

### Drug resistance phenotypic and genotypic profile

The minimum inhibitory concentrations (MICs) of different antimicrobial drugs for the four *M. brumae* strains was determined using broth microdilution panels (Sensititre RAPMYCOI and SLOMYCOI) (Supplementary Table S6). The test results were identical for the four strains and demonstrated resistance to p-aminosalicylic acid, cefepime, isoniazid (≤ 10μg/mL antimicrobial concentrations), and rifampin and intermediate susceptibility to ceftriaxone. The four *M. brumae* strains proved susceptibility to amikacin, amoxicillin/clavulanic acid 2:1, capreomycin, cefoxitin, ciprofloxacin, clarithromycin, cycloserine, doxycycline, ethambutol, ethionamide, imipenem, isoniazid (≥ 10 μg/mL antimicrobial concentration), kanamycin, linezolid, minocycline, moxifloxacin, rifabutin, streptomycin, sulfamethoxazole, tigecycline and tobramycin.

To explain the antibiotic resistance profile of *M. brumae*, drug-resistance-related mutations for rifampicin (RIF) and isoniazid (see Methods) were analyzed. Regarding RIF resistance, we did not find any of the 90 reported mutations in the *rpoB* gene associated with resistance in *M. tuberculosis*. Similarly, for isoniazid resistance, 9 reported mutations that included the *katG* gene and promoter mutations upstream of the *fabG1-inhA* operon were not found in *M. brumae*. However, the *M. brumae* sequence held several mutations in these regions compared to the sequence of *M. tuberculosis* susceptible strains (Supplementary Figure S2), indicating that the drugresistance associated mutations could be potentially located in other parts of these genes.

### Virulence related genes

For the presence or absence of M. tuberculosis virulence genetic factors in *M. brumae* and in *M. bovis* BCG Connaught, we focused on a list of M. tuberculosis virulence related genes which include PE/PPE proteins, ESX export systems, Mce proteins, MmpL proteins and proteins from the Two-Component System (TCS) among others (Supplementary Table S2). We found that *M. brumae* only held 57 out of 324 M. tuberculosis virulence-associated genes with a range of protein identity from 71.0% to 98.7% covering betwen 92.7% and 100% of the *M. tuberculosis* gene sequence (Table 2). From these, 56 genes were also shared with the *M. bovis* BCG strain (Figure 5A). Only the *phoT* protein, involved in the import of inorganic phosphate across the membrane, was found in *M. brumae* but not in *M. bovis* BCG Connaught, due to a frameshift mutation in BCG that alters the aminoacid sequence from position 156 and likely alters *phoT* function (Collins et al., 2003). We did not detect in *M. brumae* genes from PE/PPE, *Mce* or *MmpL* families with more than 70% protein identity to *M. tuberculosis* or *M. bovis* BCG, which was consist with the lack of these genes in the annotation performed with Prokka. However, *M. brumae* PGAP annotation predicted the presence of 9 *MmpL*, 15 *Mce* and 11 PE-PPE family genes. *MmpL* and *Mce* genes showed high protein identity to the ones annotated in previously published *M. brumae* contigs and in the closely related species *M. fallax*. However, the 11 PE-PPE family genes showed very low protein similarity or coverage to both. However, because the synteny of PE/PPE within ESX regions was consistent with the gene organization known for other mycobacteria, we kept them in the annotation. Although not all ESX clusters include *M. tuberculosis* virulence genes, we next analyzed in detail the presence or absence of the five ESX clusters in *M. brumae* and we found that *M. brumae* only had ESX-3 and ESX-4 gene clusters (Supplementary Figure S3), although one gene in ESX-3 is annotated as putative pseudogene (eccD4), indicated in Supplementary Table S4. *M. brumae* does not contain the virulence related ESX-1, which is in accordance with its absence in all the species clustering together in the phylogeny: *M. fallax, M. chitae* and *M. confluentis* (Figure 4). However, although ESX-2 was not complete in *M. brumae*, we found 6 genes that could be homologous to half of the genes in the ESX-2 *M. tuberculosis* H37Rv cluster (Rv3884c to Rv3889c).

**Figure 5.**
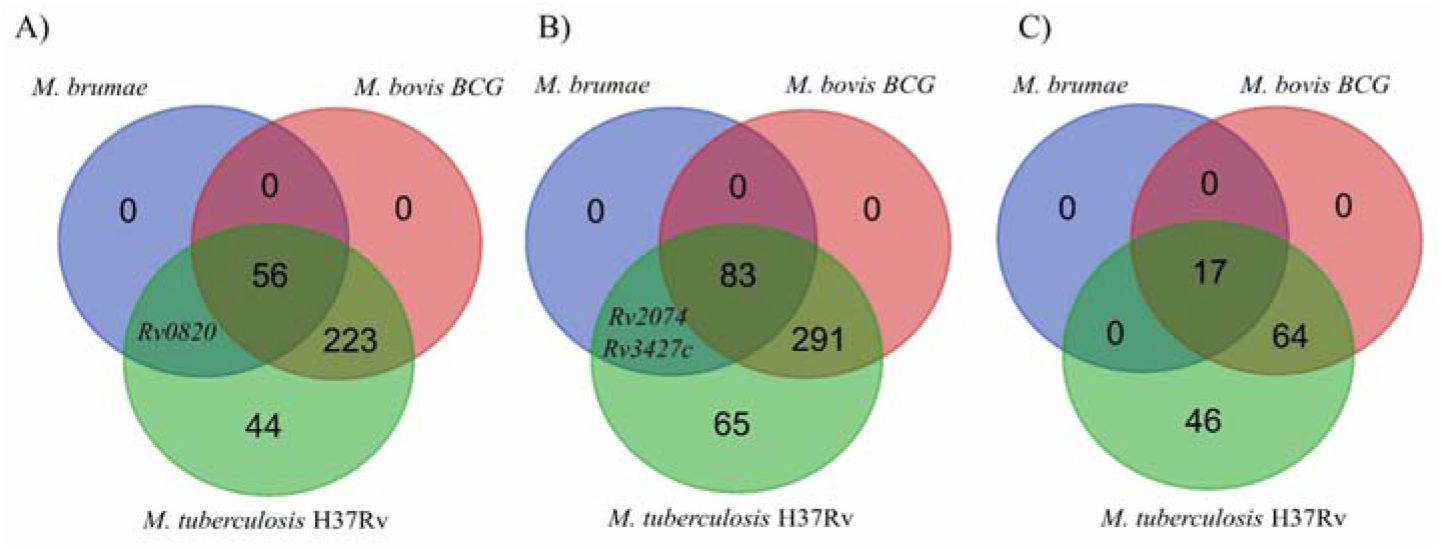
Venn diagram showing the comparison of the presence/absence and abundance of *M. tuberculosis* virulence-associated genes (A), antigens recognized by T cells (B) and antigens recognized by B cells (C) in the *M. bovis* BCG Connaught and *M. brumae* genomes.

**Table 2.**
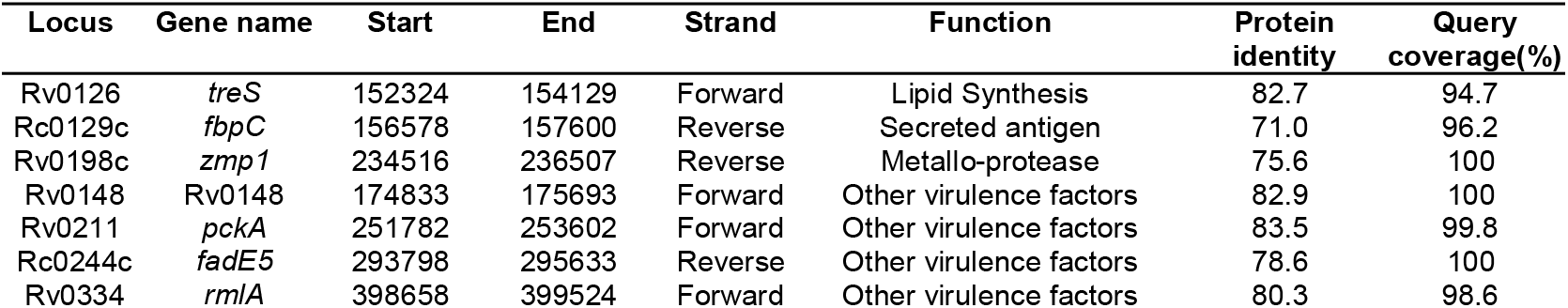
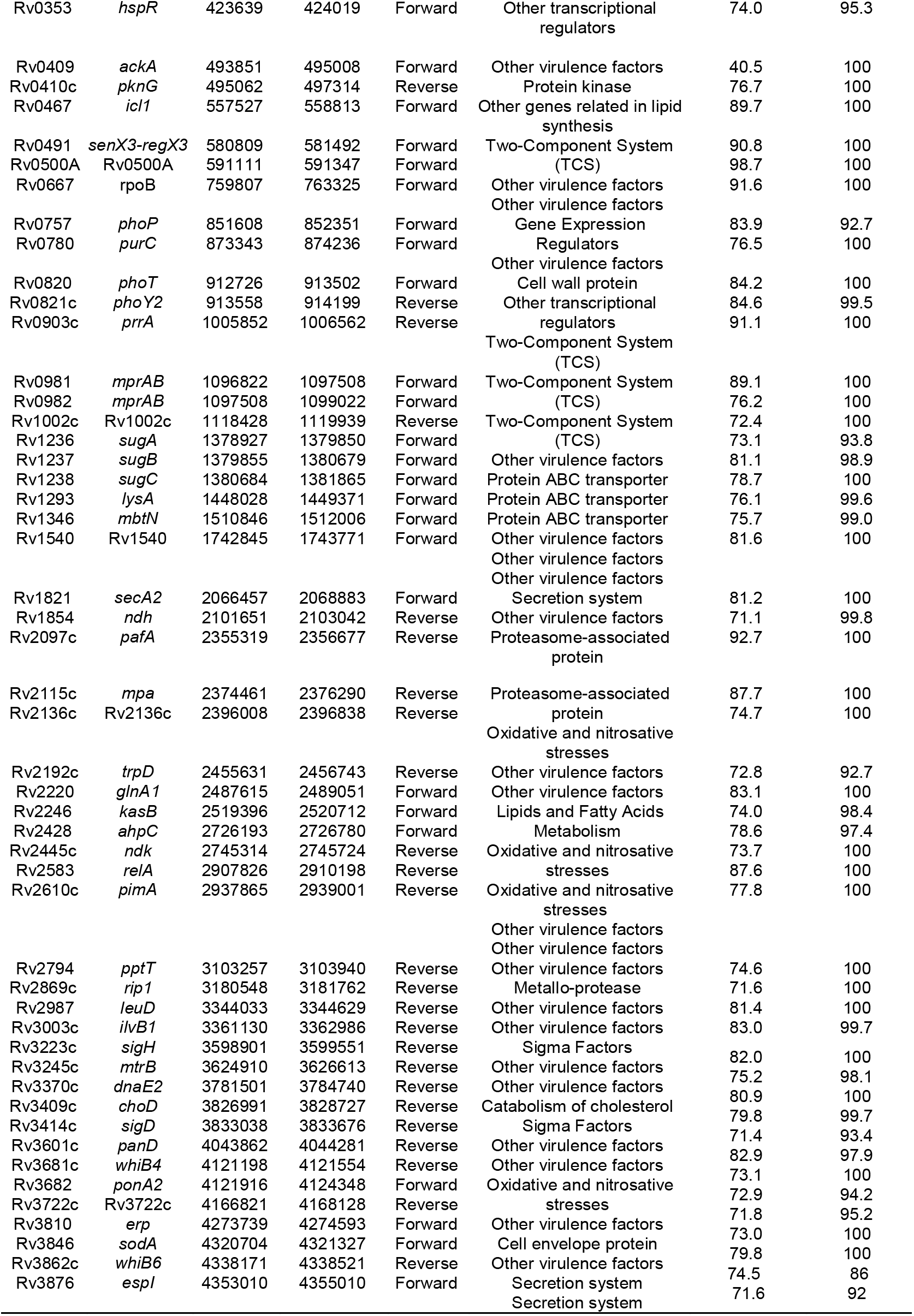
*M. tuberculosis* virulence genes with high similarity in *M. brumae*. List of the virulence-associated genes from *M. tuberculosis* H37Rv found with high similarity in the newly sequenced *M. brumae* ATCC 51384^⊤^ strain. The respective *M. tuberculosis* Rv code, gene name, orientation, position within the genome, predicted function, protein identity compared to *M. tuberculosis* and query coverage are showed.

### Comparison of the immunogenic capacity

We next analyzed the immunogenic capacity of *M. brumae* as a potential immunotherapy agent. Immunity to infections caused by mycobacteria species such as Mycobacterium tuberculosis depends mainly on T lymphocytes. For the in-silico analysis of the antigens in *M. brumae*, experimentally proved human T cell epitopes were obtained and the corresponding gene for each epitope was assessed (see Methods), obtaining a list of 442 non-redundant antigens, including the ESAT-6 gene cluster as potent immunogenic and virulence-associated regions. The presence or absence of each gene was evaluated and compared in the *M. bovis* BCG Connaught and *M. brumae* ATCC 51384T strains. As a result, we found that *M. bovis* BCG strain shared 374 out of the 441 immune-related genes with *M. tuberculosis*, while the *M. brumae* genome only had 85 genes (Supplementary Table S2), from which 83 genes were shared with the *M. bovis* BCG and the genes Rv2074 (a possible pyridoxamine 5’-phosphate oxidase) and Rv3427c, a possible transposase, were only found in *M. brumae* (Figure 5B). Interestingly, there were no ESAT-6 antigens among the genes found in *M. brumae*, which correlates with its non-virulent phenotype and shows that the immunogenic activity in *M. brumae* could be produced by other genes. B cells and humoral immunity can modulate the immune response to different pathogens, including *M. tuberculosis*. To explore other mechanisms for immunity that could modulate the immune response, we also analyzed the presence of B cell antigens in *M. brumae*. From a list of 127 non-redundant genes related to the immunogenic activity mediated by B cells, the *M. bovis* BCG strain shared 81 genes with *M. tuberculosis*, while the *M. brumae* genome only contained 17 genes (Supplementary Table S2), all of them shared with *M. bovis* BCG (Figure 5C).

### Cell wall composition

To compare the cell wall composition at the genetic level, we assessed the presence in *M. brumae* of 179 genes related to the mycobacterial cell wall (see Methods) and associated with the synthesis and transport of mycolic acids, PDIM, PGL, PIM, PL and the TDM and TMM glycolipids. As expected, the *M. brumae* genome contained genes for the synthesis and transport of α-mycolic acids and for the PIM, TDM and TMM production, and lacked PDIM and PGL associated genes (Supplementary Table S2). Interestingly, among the 96 genes related to mycolic acid production, we only found in *M. brumae* 31 mycolic-acids-associated genes, associated to general fatty acid biosynthesis functions. From the 83 total assessed genes related to the PIM, TDM, TMM and other PL production, we detected 13 genes in *M. brumae* (Rv0126, Rv0129c, Rv0982, Rv1166, Rv1236-Rv1238, Rv1564c, Rv2188, Rv2610c, Rv2869c, Rv3264c and Rv3793).

### Intraspecific variability

To assess the intraspecific variation among the different *M. brumae* strains, we analyzed the genomes of three isolates (CR103, CR142 and CR269 strains). The sequences from the three different *M. brumae* strains were uploaded to the ENA-EBI database under the BioProject code PRJEB52012 and with accession numbers ERR9463983 (CR103), ERR9463984 (CR142) and ERR9463985 (CR269). Intraspecies variability was assessed by calling SNPs and deletions compared to our newly sequenced

*M. brumae* ATCC 51384T complete genome using two different pipelines (see Methods). Mapping qualities were sufficient to study variability in SNPs and regions of difference (Supplementary Table S7). Both pipelines predicted similar number of SNPs prior to filtering, and exactly the same number of SNPs after filtering for the SNPs found in non-repetitive regions. All variable SNPs located in repeated regions that were discarded are shown in Supplementary Table S8. Strains CR103, CR142 and CR269 differed from the reference genome in 5, 3 and 6 high confidence SNPs respectively. The majority (5 of 8) of the SNPs among *M. brumae* strains were present in coding regions (Table 3). Compared to *M. brumae* ATCC 51843T reference, the three *M. brumae* isolates presented a missense variant related to the aminopeptidase N and another SNP located in a pseudogene. The CR103 strain presented only one unique SNP in a hypothetical protein (missense variant). The CR142 strain showed one unique SNP related to the propionyl-CoA--succinate CoA transferase (synonymous variant). The CR269 harbored two unique SNPs in intergenic regions which could potentially modify genes related to the glycerol-3-phosphate dehydrogenase/oxidase and the aquaporin Z.

**Table 3.**
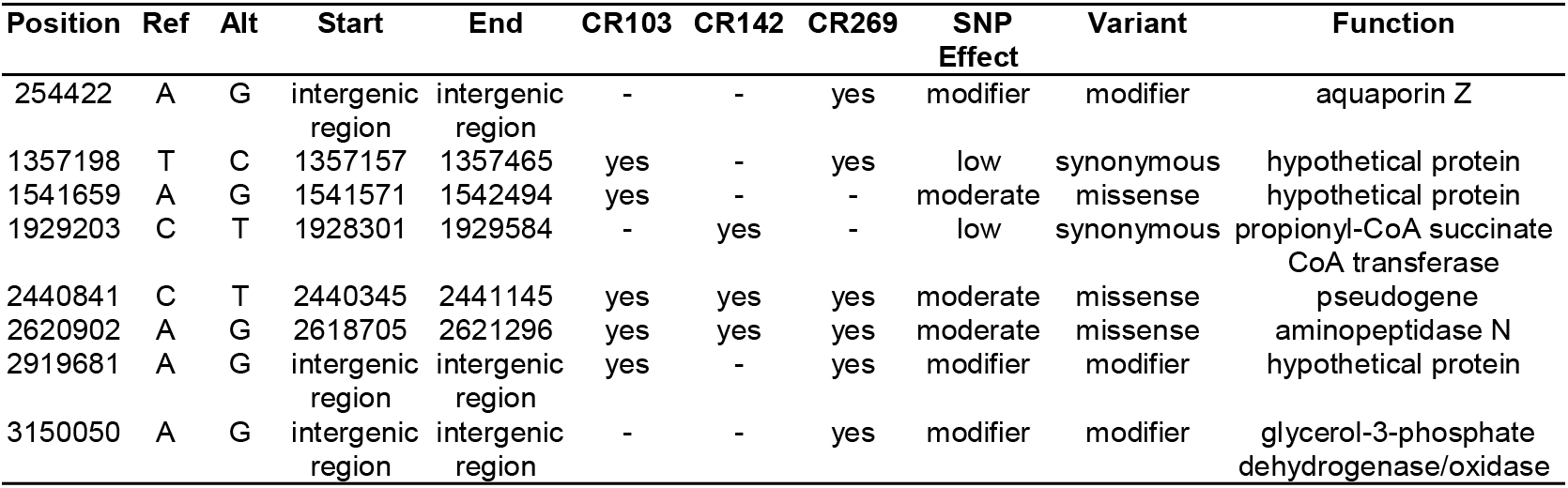
Predicted impact and characteristics of the SNPs present in the M. brumae strains compared to the *M. brumae* reference genome. SNPs found with both variant calling pipelines and outside of repetitive regions (see Methods). The table shows the position of the SNP within the reference genome, the reference base and the respective SNP found, the CDS start and end positions, the presence (“yes”) or absence (“-”) of the SNP in each *M. brumae* strain and the predicted impact of the mutation by the SnpEff software. The protein function is decided for the involved CDS or the following gene in intergenic regions.

We also examined if genes from the reference ATCC 51384T strain were absent in any of the three *M. brumae* genomes, by analyzing the coverage of Illumina reads in each position (see Methods). We found up to six deleted regions. Only one deletion involved two genes (annotated as infB and YlxR family protein, while the others were involved in the translation of only one protein product. Other affected genes were a transglycosylase, a C40 family peptidase, apa (Rv1860), and hypothetical proteins (Table 4). Four regions were absent in all three strains compared with the reference genome. CR103 and CR142 lacked apa gene and two hypothetical proteins were absent in CR269 (Table 4).

**Table 4.** Genes affected by deletions in the different *M. brumae* strains compared to the *M. brumae* reference genome. CDS ID indicates the PGAP identity code for genes affected by deletions. The product obtained for each gene was predicted by PGAP and confirmed by blastp in *M. tuberculosis* H37Rv.

| Gene annotation |  | Percentage deleted (%) |  |  |
| --- | --- | --- | --- | --- |
| CDS ID | Product | CR103 | CR142 | CR269 |
| pgaptmp_000208 | Transglycosylase family protein | 37.02 | 37.16 | 32.92 |
| pgaptmp_001051 | Hypothetical protein | 29.05 | 29.05 | - |
| pgaptmp_001263 | C40 family peptidase | 37.79 | 32.34 | 38.94 |
| pgaptmp_001306 | *apa* (Rv1860) | - | - | 25.30 |
| pgaptmp_002029 | *infB* | 42.92 | 40.31 | 44.99 |
| pgaptmp_002030 | YlxR family protein | 25.08 | 25.08 | 27.65 |
| pgaptmp_003794 | Hypothetical protein | - | 26.16 | - |

## Discussion

In this study, we assembly and compare the genome of the *M. brumae* type strain with *M. bovis* BCG and *M. tuberculosis* to precisely unravel the peculiar characteristics shown phenotypically by this species: its non-pathogenicity and high immunostimulatory ability. We also compare some genomic features of *M. brumae* with *M. fallax*, a fast-growing mycobacterial species that was initially isolated from water samples. M. fallax shares phenotypic characteristics with *M. brumae* and in initial phylogenetic comparisons appeared as the closest species to *M. brumae* (Luquin et al., 1993; Noguera-Ortega, 2016a). The phylogenetic position of *M. brumae* is in accordance with the microbiological features observed (Bach-Griera et al., 2020) as a non-pathogenic rapidly-growing mycobacteria. Genome-based phylogenetic reconstruction confirms the position of *M. brumae* within *Mycobacteriaceae*, and closely related to *M. fallax*, which was shown in previous analysis with limited data based on 16S rRNA sequences (Noguera-Ortega et al., 2016b).

We confirmed the small size of *M. brumae* genome. Although it is accepted that the size of the genome of nonpathogenic mycobacteria is longer that those of opportunistic and obligated pathogenic mycobacteria (Rahman et al., 2014; Jia et al., 2021), M. brumae genome is of similar size than *M. tuberculosis* genome comparable only to their related Mycobacterium species such as *M. fallax*. In fact, *M. brumae* genome is, as far we know, the second smallest genome among mycobacteria followed by the pathogenic Mycobacterium leprae with a genome size of 3.3 Mb (Rahman et al., 2014; Matsumoto et al., 2019). Our results are highly congruent with the draft genome obtained with an Illumina MiSeq which had a length of 4,026,006 bp and a mean GC ratio of 69.1% (D’Auria et al., 2016). The percentage of GC content in *M. brumae* genome is in accordance with other RGMs genomes (between 66.0 to 69.0%) and higher than for slow-growing mycobacteria; only *M. chelonae-M. abscessus* group contain a percentage between 63.9 to 64.1% of GC contents among RGMs (Sassi and Drancourt, 2014). Despite sharing a genome size with pathogenic mycobacteria, the pattern of gene categories is different between *M. brumae* and *M. tuberculosis* or *M. bovis* BCG. A lower percentage of genes related to defense mechanisms and lipid metabolism, among others, is found in *M. brumae* in comparison to *M. tuberculosis* genome, which correlates with its non-pathogenicity (Gago et al., 2018).

The mycobacterial Type VII secretion system (Type VII) (ESX1-5) is specialized for the secretion of various protein substrates across the complex cell envelope, and they have evolved through duplication leading to a variety of ESX clusters across the mycobacterial genus (Newton-Foot et al., 2016). *M. tuberculosis* and some nonpathogenic Mycobacterium such as *Mycobacterium paragordonae* (Kim et al., 2019) harbors five ESX clusters, other non-pathogenic mycobacteria such as *Mycobacterium smegmatis, Mycobacterium flavescens, M. phlei, Mycobacterium gilvum, Mycobacterium vaccae*, or *Mycobacterium gordonae* only three (Gcebe et al., 2016; Newton-Foot et al., 2016). Here we show that, like *M. abcessus* and *M. massiliensse* (Gcebe et al., 2016), *M. brumae* genome harbors only ESX-4 and ESX-3 and lacks other ESX clusters. The ESX-1 system has been implicated in virulence, and its deletion has been related to the attenuation of the pathogenic mycobacteria as in the case of *M. bovis* BCG (Hsu et al., 2003; Pym et al., 2003). However, its presence throughout the *Mycobacterium* genus, including non-pathogenic and fast-growing organisms (Gcebe et al., 2016) suggests that the primary function of this gene cluster is not the virulence, and that the virulence-associated function could have evolved more recently in pathogenic organisms. PE/PPE family proteins represent about 10% of the total *M. tuberculosis* genome and they are expressed by *M. tuberculosis* upon infection of macrophages and play critical roles in virulence, antigenic diversity, and modulation of the host immune response (Mukhopadhyay and Balaji, 2011; Fishbein et al., 2015). Similar to what is found for *M. fallax*, very few PE/PPE family genes are annotated in *M. brumae* genome, and sequence identity from the few annotated PE/PPE genes is very low compared to *M. tuberculosis* and *M. fallax*. The distant nature and the low number of these genes in *M. brumae* and in *M. fallax* is consistent with the fact that these genes are more abundant in slow-growing mycobacteria (Qian et al., 2020) (Gey van Pittius et al., 2006).

Mce proteins are lipid/sterol transporters (Cantrell et al., 2013) that play an important role interfering the host cell signaling modulation (Fenn et al., 2019) and they are implicated in the entry and survival inside macrophages (Saini et al., 2008; Fenn et al., 2019). We could only detect three mce cluster *M. brumae* genome but showing very low sequence similarity with *M. tuberculosis mce* genes, which impedes to accurately hypothesize its orthology with specific *mce* clusters in *M. tuberculosis*. The number of mce operons present in the genome has been related to pathogenicity in actinomycetes: both in mycobacteria and nocardia. *M. abscessus* contains seven mce operons (mce1-7) while *M. smegmatis* only four (Ripoll et al., 2009). There are six mce operons in Nocardia farcinica, one of the agents causing nocardiosis, whereas *Streptomyces avermitilis* and *Streptomyces coelicolor*, both nonpathogenic soil bacteria, each have only one copy of the mce operon (Ishikawa et al., 2004). In a recent study, Bachmann et al., 2020 found that the mce1 operon is highly variable across all five mycobacterial sub-genera. The study indicates that the sequence similarity of proteins encoded by the mce1 genes between *M. tuberculosis* and *M. brumae* was very low, which confirms our results. Although there is yet no described genetic or phenotypic markers whose appearance correlates with nontuberculous mycobacterial virulence (Falkinham, 2009), the absence of virulence genes, such as ESX-1, and most PE/PPE can explain the results previously obtained in in vitro and in vivo models, demonstrating the safety and non-toxicity of M. brumae (summarized in Figure 1) (Noguera-Ortega et al., 2016b, 2020; Bach-Griera et al., 2020). *M. brumae*, as most of the more than 190 mycobacterial species that have been identified, have not been linked with human, animal, or plant disease. Furthermore, the absence of ESX-1 secretion system in *M. brumae* is interesting since this chromosomal region is involved in a unique conjugation mechanism to some RGM, as has been demonstrated for *M. smegmatis*, potentially enabling saprophytic mycobacteria to change niche and evolve to opportunistic or specialized persistent pathogen (Coros et al., 2008; Gray and Derbyshire, 2018).

Previous studies related to resistance to the main anti-tuberculosis drugs in NTMs indicate that it is mainly due to specific mutations in genes that encode their target or the enzymes that activate them, as well as their promoter regions (Alcaide et al., 2017; Bakuła et al., 2018). Resistance to rifampin in M. tuberculosis is conferred by mutations mainly in the rpoB gene, and similar mutations in the same cluster have been described for *M. leprae* and some NTM species such as *Mycobacterium kansasii, M. smegmatis, M. avium* complex and *Mycobacterium ulcerans* (Brown-Elliott et al., 2012; van Ingen et al., 2012; Alcaide et al., 2017). *M. brumae* lacks point mutations within RIF-resistance determining region (RRDR) of the *rpoB* gene but has mutations in adjoining areas of the region. Similarly, in some resistant isolates of *M. avium* no internal mutations were found in rpoB but in a downstream region of this gene. This fact together with the mycobacterial cell wall composition, which has a role in the intrinsic resistance to rifampin (Brown-Elliott et al., 2012; Huh et al., 2019), could explain the resistance to rifampin observed in *M. brumae*. Regarding isoniazid resistance, often associated with a loss of catalase-peroxidase activity, point mutations in the sequence of *katG* gene (responsible for activating the drug) have been described in *M. tuberculosis* and some NTMs such as *M. smegmatis* or *M. kansasii* (Brown-Elliott et al., 2012; Alcaide et al., 2017; Iwao and Nakata, 2018). In addition, resistance to isoniazid has been described in *M. smegmatis, M. bovis* BCG and *M. tuberculosis* due to overexpression or mutation of the inhA structural gene or its promoter, which are involved in the synthesis of the bacterial wall (Larsen et al., 2002; Brown-Elliott et al., 2012; Alcaide et al., 2017). *M. brumae* strains lack point mutations in the most relevant positions described for isoniazid. However, the low cell wall permeability of mycobacteria and the different efflux pump systems or porin channels, could also play a crucial role in intrinsic drug resistance in NTMs such as M. fortuitum or *M. smegmatis* (van Ingen et al., 2012; Bakuła et al., 2018) and could explain the intrinsic resistance to isoniazid found in *M. brumae*. Finally, similar to isolates of *M. fortuitum* and *M. avium* complex (Cheng et al., 2017), *M. brumae* is also resistant to p-aminosalicylic acid, although the mutations responsible for resistance have not been specifically identified (Yang et al., 2022). Thus, due to the differences in drug resistance genetic determinants among *Mycobacterium* species, additional efforts are needed to determine drug susceptibility by molecular approaches in NTM as *M. brumae*.

The comparative analysis of immunogenic T cell and B cell epitopes highlights that *M. brumae* is sharing very few antigens with BCG and *M. tuberculosis* and it can still trigger an immune response to similar levels than BCG in preclinical studies related to bladder cancer immunotherapy (summarized in Figure 6). *M. brumae* might have other immunogenic epitopes that are not described in *M. tuberculosis*, and finding those would provide a better knowledge of *M. brumae* immunogenic profile. Interestingly, some of the shared genes are critical immunogenic heat-shock proteins (groEL, groES or dnaK) involved in the immunostimulatory ability of BCG (Qazi et al., 2005). Heatshock proteins have been the focus of numerous studies as potential immunotherapy tools in different types of cancer (Das et al., 2019). In fact, purified heat-shock proteins from mycobacteria species or BCG overexpressing diverse heat-shock proteins have been studied as adjuvants for the treatment of different cancers with some success (reviewed in (Noguera-Ortega et al., 2020). Therefore, the high *M. brumae* immune potential could be related to a higher expression of antigens or the presence of alternative immunogenic molecules. Another source of immunogenic molecules is the mycobacterial cell wall. The cell wall biosynthesis genes found in the *M. brumae* genome is in accordance with the cell wall composition described previously for this species (Guallar-Garrido et al., 2021) (Supplementary Figure S4). The hallmark of mycobacteria is their unique abundance of lipids and glycolipids in their cell wall (Chiaradia et al., 2017; Minnikin and Brennan, 2020), which include the exceptionally long chain fatty acids (mycolic acids, MA). Mycolic acids are useful taxonomic markers, and they are related to host-mycobacteria interaction. *M. brumae* is one of the few species that only contains α-mycolic acids (type I) (Luquin et al., 1993) (Supplementary Figure S4). Like most mycobacteria species, *M. brumae* contains TMM, TDM, PIM and PL (Guallar-Garrido et al., 2021). However, hydrophilic glycopeptidolipids (GPLs) or lipooligosaccharides (LOS), present in other NTM are absent in *M. brumae*. Unlike *M. bovis* BCG, *M. brumae* does not contain lipids related to virulence like highly hydrophobic PDIM or their related PGLs, or sulfolipid (SL) and di-acyl- and poly-acyl trehaloses (DAT, PAT) present in the M. tuberculosis cell wall (Minnikin et al., 2002) (Supplementary Figure S4).

**Figure 6.**
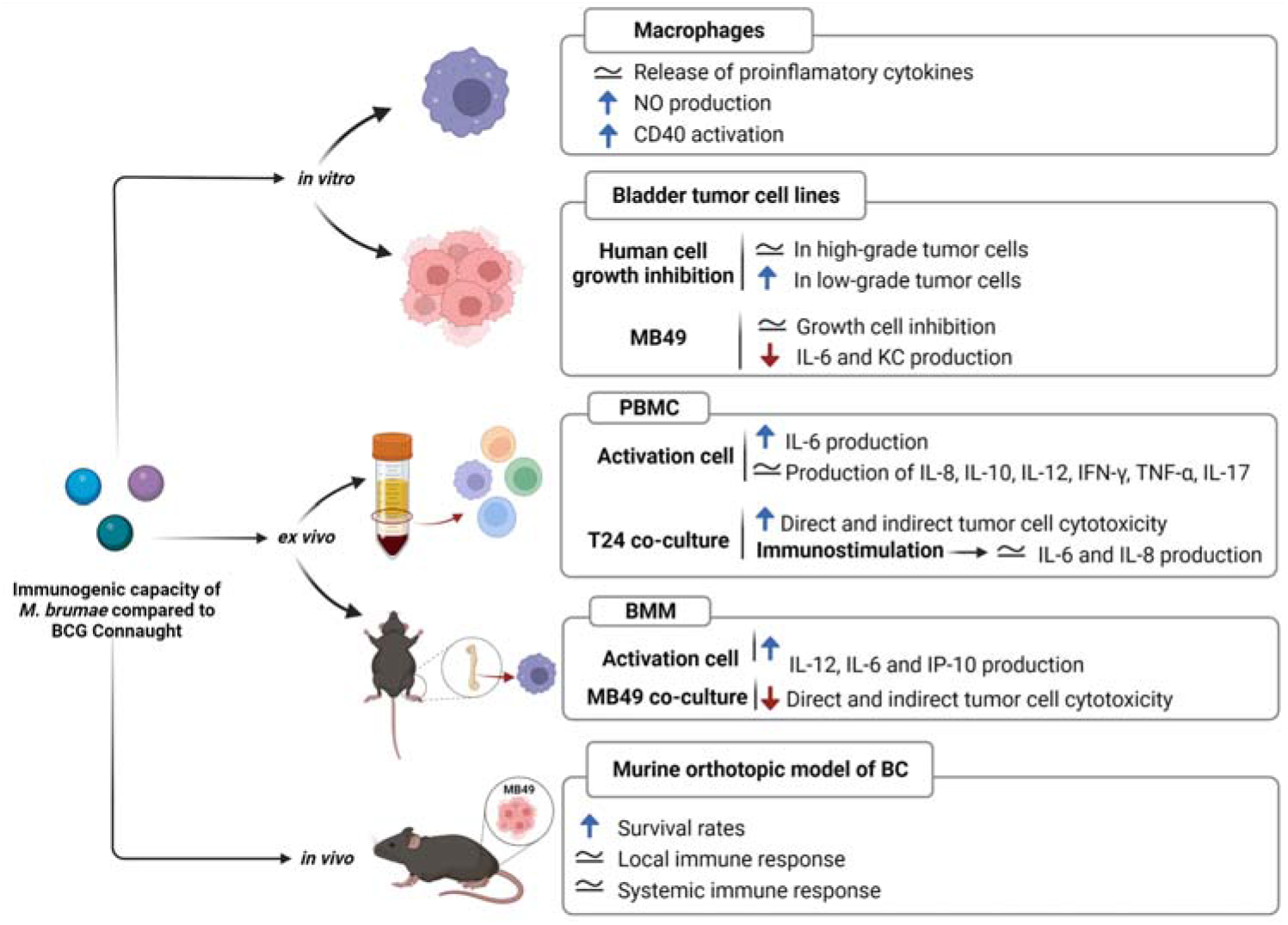
Immunogenic capacity of *M. brumae* compared *to M. bovis* BCG Connaught. Summary of the results obtained in studies treating with both mycobacteria J774 macrophages cell line (Noguera-Ortega et al., 2016b), bladder cancer cell lines (Noguera-Ortega et al., 2016a), human peripheral mononuclear cells (Noguera-Ortega et al., 2016b), and murine bone-marrow macrophages (unpublished)); and after treating intravesically with each mycobacteria bladder cancer tumor-bearing mice (Noguera-Ortega et al., 2016a, 2016b, 2018)). Blue rising arrow indicates higher than *M. bovis* BCG Connaught, red falling arrow indicates lower than *M. bovis* BCG Connaught and equals symbol indicates similar to *M. bovis* BCG Connaught.

Apart from the reference strain, other three *M. brumae* strains are available. The type *M. brumae* strain (ATCC 51384T initially named as CR270), and CR103 and CR142 strains were isolated from water samples taken from the Llobregat River in Spain between 1983 and 1987, and CR269 from soil between 1989 and 1991 (Luquin et al., 1993). Despite coming from slighty different environmental sources due to a different timeframe, all four strains showed similar *in vitro* genotypes and high genomic similarity. Phenotypically, they showed the same colony morphology when grown in solid media (Supplementary Figure S5), similar pattern of lipids and glycolipids on the cell wall and the same susceptibility pattern to a wide range of antibiotics. At the genome level, they showed a maximum of six SNPs and seven regions of difference, which is a very low genomic diversity for different environmental strains. Other mycobacteria, such as *M. abscessus* have shown very low genetic diversity across wide temporal and geographical scales too. Person to person transmission has been refuted to be the reason for the low genetic diversity in *M. abscessus*, and a lower mutation rate due to differences in a repair mechanism has been proposed as the most probable explanation (Commins et al., 2022).

In summary, the *M. brumae* diversity analysis demonstrated that *M. brumae* is genetically less diverse than expected from the diverse origin of the strains compared. The gene composition reported reveals that *M. brumae* interacts differently to host immune system compared to host-specialized mycobacterial groups, such as *M. bovis* BCG or M. tuberculosis. Our research not only reveals the variety genomic features among nontuberculous mycobacteria species, but it also provides an insight into finding critical antigenic molecules that could be the basis for designing bacteria-based therapies to combat cancer or other immune-based disorders.

## Acknowledgments

We acknowledge the access to Centro de Investigaciones Príncipe Felipe (CIPF) scientific computing core facility in Valencia.

