## Supplementary material for "Genomic analysis of *Mycobacterium brumae* sustains its nonpathogenic and immunogenic phenotype": Data Sheet 1.PDF

**Supplementary Figure S3.** Phylogenetic relationship of the available *M. brumae* genomic sequences. A) Maximum likelihood phylogenetic tree based on an alignment of the core-genome with a length of 293,792 bp and including *M. fallax* as outgroup. GTR model was used with bootstrap confidence of 1000 replicates. B) Zoom-in to the *M. brumae* cluster of the previous tree showed in A. Red dots indicate genomic sequences of the type strain (same as ATCC 51384).

A)

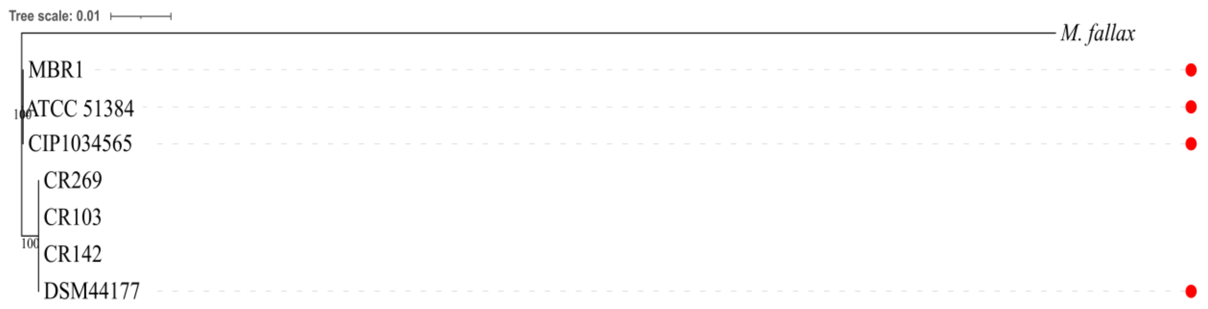


B)

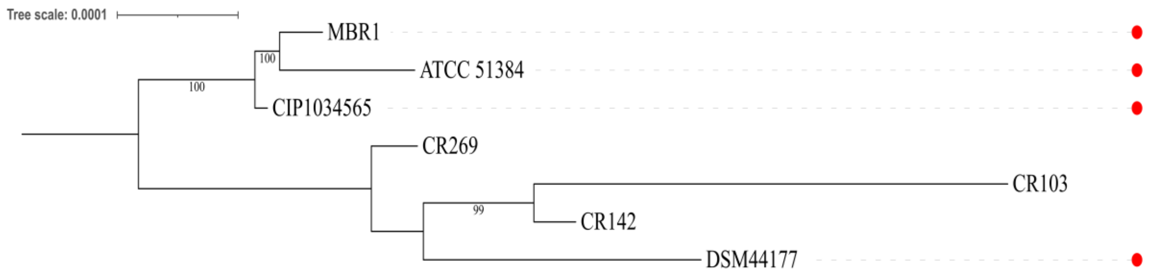
