## Supplementary material for "Genomic analysis of *Mycobacterium brumae* sustains its nonpathogenic and immunogenic phenotype": Data Sheet 2.PDF

**Supplementary Figure S2.** Protein alignments for the analysis of drug-resistance mutations. Query represents *M. tuberculosis* H37Rv and Sbjct represents *M. brumae*. All *M. brumae* mutations are indicated in the alignment. **A)** RpoB protein alignment for rifampicin resistance analysis. **B)** KatG protein alignment for isoniazid resistance analysis.

A)

```
Score            Expect Method                       Identities         Positives          Gaps
2151 bits(5574)  0.0    Compositional matrix adjust. 1059/1179(90%)     1113/1179(94%)     18/1179(1%)

Query  1     LADSRQSKTAASPSPSRPQSSSNNSVPGAPNRVSFAKLREPLEVPGLLDVQTDSFEWLIG  60
             LA SRQSK+ A+           NSVPGAPNR+SFAKLREPLEVPGLLDVQTDSFEWLIG
Sbjct  7     LAVSRQSKSVAT-----------NSVPGAPNRISFAKLREPLEVPGLLDVQTDSFEWLIG  55

Query  61    SPRWRESAAERGDV-NPVGGLEEVLYELSPIEDFSGSMSLSFSDPRFDDVKAPVDECKDK  119
             S +WR SA+ RGD+ NP GGLEEVL ELSPIEDFSGSMSLSFSDPRFD+VKAPV ECKDK
Sbjct  56    SDKWRASASARGDIANPKGGLEEVLDELSPIEDFSGSMSLSFSDPRFDEVKAPVAECKDK  115

Query  120   DMTYAAPLFVTAEFINNNTGEIKSQTVFMGDFPMMTEKGTFIINGTERVVVSQLVRSPGV  179
             DMTYAAPLFVTAEFINNNTGEIKSQTVFMGDFPMMTEKGTFIINGTERVVVSQLVRSPGV
Sbjct  116   DMTYAAPLFVTAEFINNNTGEIKSQTVFMGDFPMMTEKGTFIINGTERVVVSQLVRSPGV  175

Query  180   YFDETIDKSTDKTLHSVKVIPSRGAWLEFDVDKRDTVGVRIDRKRRQPVTVLLKALGWTS  239
             YFDE+IDKST+KTLHSVKVIP RGAWLEFDVDKRDTVGVRIDRKRRQPVTVLLKALGWTS
Sbjct  176   YFDESIDKSTEKTLHSVKVIPGRGAWLEFDVDKRDTVGVRIDRKRRQPVTVLLKALGWTS  235

Query  240   EQIVERFGFSEIMRSTLEKDNTVGTDEALLDIYRKLRPGEPPTKESAQTLLENLFFKEKR  299
             EQI ERFGFSEIM  TLEKD T G DEALLDIYRKLRPGEPPTKESAQTLLENLFFKEKR
Sbjct  236   EQIRERFGFSEIMMGTLEKDPTAGPDEALLDIYRKLRPGEPPTKESAQTLLENLFFKEKR  295

Query  300   YDLARVGRYKVNKKLGLHVGEPI--TSSTLTEEDVVATIEYLVRLHEGQTTMTVPGGVEV  357
             YDLARVGRYKVNKKLGL++ +P+  +S+TLTEED+VATIEYLVRLHEGQTTMT PGG EV
Sbjct  296   YDLARVGRYKVNKKLGLNLDKPVNTSSNTLTEEDIVATIEYLVRLHEGQTTMTAPGGSEV  355

Query  358   PVETDDIDHFGNRRLRTVGELIQNQIRVGMSRMERVVRERMTTQDVEAITPQTLINIRPV  417
             PVE DDIDHFGNRRLRTVGELIQNQIRVG+SRMERVVRERMTTQDVEAITPQTLINIRPV
Sbjct  356   PVEVDDIDHFGNRRLRTVGELIQNQIRVGLSRMERVVRERMTTQDVEAITPQTLINIRPV  415

Query  418   VAAIKEFFGTSQLSQFMDQNNPLSGLTHKRRLSALGPGGLSRERAGLEVRDVHPSHYGRM  477
             VAAIKEFFGTSQLSQFMDQNNPLSGLTHKRRLSALGPGGLSRERAGLEVRDVH SHYGRM
Sbjct  416   VAAIKEFFGTSQLSQFMDQNNPLSGLTHKRRLSALGPGGLSRERAGLEVRDVHSSHYGRM  475

Query  478   CPIETPEGPNIGLIGSLSVYARVNPFGFIETPYRKVVDGVVSDEIVYLTADEEDRHVVAQ  537
             CPIETPEGPNIGLIGSLSVYARVNPFGFIETPYRKV+DGV++D+I YLTADEEDRHVVAQ
Sbjct  476   CPIETPEGPNIGLIGSLSVYARVNPFGFIETPYRKVIDGVITDQIDYLTADEEDRHVVAQ  535

Query  538   ANSPI-DADGR--FVEPRVLVRRKAGEVEYVPSSEVDYMDVSPRQMVSVATAMIPFLEHD  594
             ANSP  + DG   F + RVLVRRK GEVE+V ++EVDYMDVSPRQMVSVATAMIPFLEHD
Sbjct  536   ANSPTAEGDGGLVFTDDRVLVRRKGGEVEFVSAAEVDYMDVSPRQMVSVATAMIPFLEHD  595

Query  595   DANRALMGANMQRQAVPLVRSEAPLVGTGMELRAAIDAGDVVVAEESGVIEEVSADYITV  654
             DANRALMGANMQRQAVPLVRSEAPLVGTGMELRAAIDAGDVVV+E++GV+EEVSADYITV
Sbjct  596   DANRALMGANMQRQAVPLVRSEAPLVGTGMELRAAIDAGDVVVSEKAGVVEEVSADYITV  655

Query  655   MHDNGTRRTYRMRKFARSNHGTCANQCPIVDAGDRVEAGQVIADGPCTDDGEMALGKNLL  714
             M D+GTR+TYRMRKFARSNHGTCANQ PIVDAG RVE+GQV+ADGPCT++GEMALGKNLL
Sbjct  656   MADDGTRQTYRMRKFARSNHGTCANQRPIVDAGQRVESGQVVADGPCTENGEMALGKNLL  715

Query  715   VAIMPWEGHNYEDAIILSNRLVEEDVLTSIHIEEHEIDARDTKLGAEEITRDIPNISDEV  774
             VA+MPWEGHNYEDAIILS RLVEEDVLTSIHIEEHEIDARDTKLGAEEITRDIPN+SDEV
Sbjct  716   VAVMPWEGHNYEDAIILSQRLVEEDVLTSIHIEEHEIDARDTKLGAEEITRDIPNVSDEV  775

Query  775   LADLDERGIVRIGAEVRDGDILVGKVTPKGETELTPEERLLRAIFGEKAREVRDTSLKVP  834
             LADLDERGIVRIGAEVRDGDILVGKVTPKGETELTPEERLLRAIFGEKAREVRDTSLKVP
Sbjct  776   LADLDERGIVRIGAEVRDGDILVGKVTPKGETELTPEERLLRAIFGEKAREVRDTSLKVP  835

Query  835   HGESGKVIGIRVFSREDEDELPAGVNELVRVYVAQKRKISDGDKLAGRHGNKGVIGKILP  894
             HGESGKVIGIRVFSRED+DELPAGVNELVRVYVAQKRKISDGDKLAGRHGNKGVIGKILP
Sbjct  836   HGESGKVIGIRVFSREDDDELPAGVNELVRVYVAQKRKISDGDKLAGRHGNKGVIGKILP  895

Query  895   VEDMPFLADGTPVDIILNTHGVPRRMNIGQILETHLGWCAHSGWKVDAAKG-VPDWAARL  953
             +EDMPFL DGTPVDIILNTHGVPRRMNIGQILETHLGW A +GW V  A G +P+WA  L
Sbjct  896   IEDMPFLPDGTPVDIILNTHGVPRRMNIGQILETHLGWVAKAGWNVQVAGGEIPEWAGNL  955

Query  954   PDELLEAQPNAIVSTPVFDGAQEAELQGLLSCTLPNRDGDVLVDADGKAMLFDGRSGEPF  1013
             P++LL A  ++IVSTPVFDGA+E ELQGLLS TLPNRDG+V+VD DGKA LFDGRSGEPF
Sbjct  956   PEDLLSAPSDSIVSTPVFDGAREEELQGLLSSTLPNRDGEVMVDGDGKARLFDGRSGEPF  1015

Query  1014  PYPVTVGYMYIMKLHHLVDDKIHARSTGPYSMITQQPLGGKAQFGGQRFGEMECWAMQAY  1073
             PYPVTVGYMYI+KLHHLVDDKIHARSTGPYSMITQQPLGGKAQFGGQRFGEMECWAMQAY
Sbjct  1016  PYPVTVGYMYILKLHHLVDDKIHARSTGPYSMITQQPLGGKAQFGGQRFGEMECWAMQAY  1075

Query  1074  GAAYTLQELLTIKSDDTVGRVKVYEAIVKGENIPEPGIPESFKVLLKELQSLCLNVEVLS  1133
             GAAYTLQELLTIKSDDTVGRVKVYEAIVKGENIPEPGIPESFKVLLKELQSLCLNVEVLS
Sbjct  1076  GAAYTLQELLTIKSDDTVGRVKVYEAIVKGENIPEPGIPESFKVLLKELQSLCLNVEVLS  1135

Query  1134  SDGAAIELREGEDEDLERAAANLGINLSRNESASVEDLA  1172
             SDGAAIE+R+G+DEDLERAAANLGINLSRNESASVEDLA
Sbjct  1136  SDGAAIEMRDGDDEDLERAAANLGINLSRNESASVEDLA  1174
```

B)

| Score | Expect | Method | Identities | Positives | Gaps |
|---|---|---|---|---|---|
| 972 bits(2513) | 0.0 | Compositional matrix adjust. | 488/745(66%) | 576/745(77%) | 18/745(2%) |

```
Query  1    VPEQHPPITETTTGAASNGCPVVGHMKYPVEGGGNQDWWPNRLNLKVLHQNPAVADPMGA  60
            VPE+ PPI E  T    +GCP+   +K PVEGG N+DWWPN++NLK+L +NP + DP
Sbjct  1    VPEETPPIGEAQT---ESGCPM--RIKPPVEGGSNRDWWPNQVNLKILQKNPDIIDPEDE  55

Query  61   AFDYAAEVATIDVDALTRDIEEVMTTSQPWWPADYGHYGPLFIRMAWHAAGTYRIHDGRG  120
             +DY   V T+D +    D + ++T SQ WWPAD+GHYGPLF+RM+WHAAGTYR+ DGRG
Sbjct  56   GYDYRQVVQTLDFEEFQADFDALLTDSQSWWPADFGHYGPLFVRMSWHAAGTYRVQDGRG  115

Query  121  GAGGGMQRFAPLNSWPDNASLDKARRLLWPVKKKYGKKLSWADLIVFAGNCALESMGFKT  180
            GAG GMQRF PLNSWPDN  LD+ARRLLWP+KKKYG K+SWADLI +AGN A+E MGFKT
Sbjct  116  GAGRGMQRFEPLNSWPDNVLLDQARRLLWPLKKKYGNKISWADLIAYAGNRAMEHMGFKT  175

Query  181  FGFGFGRVDQWEPDE-VYWGKEATWLGDE-RYSG--KRDLENPLAAVQMGLIYVNPEGPN  236
             GF FGR D WEP+E V+WG EA WLG + RY G  +  L+NPLAA  MGLIYVNPEGP
Sbjct  176  AGFAFGRADCWEPEEDVFWGAEAEWLGSQDRYQGSDRTKLDNPLAATMMGLIYVNPEGPE  235

Query  237  GNPDPMAAAVDIRETFRRMAMNDVETAALIVGGHTFGKTHGAGPADLVGPEPEAAPLEQM  296
            G PDP+AAA+DIRETF RMAMNDVETAALIVGGHTFGKTHG G A+ +GPEP AAPL++M
Sbjct  236  GVPDPLAAAIDIRETFGRMAMNDVETAALIVGGHTFGKTHGNGDAEALGPEPAAAPLQEM  295

Query  297  GLGWKSSYGTGTGKDAITSGIEVVWTNTPTKWDNSFLEILYGYEWELTKSPAGAWQYTAK  356
            GLGWK+   TG   D + SG+EV+WT+TPTKWDNSFLEILY  EWEL KS AGA Q+  K
Sbjct  296  GLGWKNPNDTGNPNDRVGSGLEVIWTHTPTKWDNSFLEILYSNEWELFKSKAGAQQWRPK  355

Query  357  DGAGAGTIPDPFGGPGRSPTMLATDLSLRVDPIYERITRRWLEHPEELADEFAKAWYKLI  416
            D   A ++P P       P ML TDLS+R DPIY +ITRRWL+HP+ELA+EFAKAW+KL+
Sbjct  356  DNGWANSVPTPDLKGRTHPAMLTTDLSMREDPIYGKITRRWLDHPDELAEEFAKAWFKLM  415

Query  417  HRDMGPVARYLGPLVPKQTLLWQDPVPAVSHDLVGEAEIASLKSQIRASGLTVSQLVSTA  476
            HRDMGP  RYLGP VPK T +WQDPVPA + +L  +A++A+LK+ I  SGLTV QLVSTA
Sbjct  416  HRDMGPAVRYLGPFVPKDTWVWQDPVPAGNANL-SDADVAALKAAIADSGLTVPQLVSTA  474

Query  477  WAAASSFRGSDKRGGANGGRIRLQPQVGWEVNDPDGDLRKVIRTLEEIQESFNSAAPGNI  536
            W AA+S+R SD RGGANGGRIRLQPQ+GWE N+PD +L +VIR LEEIQ S        +
Sbjct  475  WKAAASYRNSDMRGGANGGRIRLQPQLGWESNEPD-ELAQVIRKLEEIQAS------SGV  527

Query  537  KVSFADLVVLGGCAAIEKAAKAAGHNITVPFTPGRTDASQEQTDVESFAVLEPKADGFRN  596
             VSFAD+VVL G   +EKAAKAAG +I VPFTPGR DA+QE TD +SF+ LEPKADGFRN
Sbjct  528  NVSFADVVVLAGNVGVEKAAKAAGFDIDVPFTPGRGDATQEMTDADSFSYLEPKADGFRN  587

Query  597  YLGKGNPLPAEYMLLDKANLLTLSAPEMTVLVGGLRVLGANYKRLPLGVFTEASESLTND  656
            Y GKG  LPAEY L+D+AN L LS PEMTVLVGGLR L AN+    LGV TE   +LT D
Sbjct  588  YAGKGLNLPAEYHLIDRANQLNLSGPEMTVLVGGLRALEANHGGSKLGVLTERPGALTTD  647

Query  657  FFVNLLDMGITWEPSPADDGTYQGKD-GSGKVKWTGSRVDLVFGSNSELRALVEVYGADD  715
            FFV++ DMG+ W PS ADDGTY G D  +G+ K+T SRVDL+FGSNS+LRAL EVY ADD
Sbjct  648  FFVSICDMGLKWAPSSADDGTYVGSDRATGEPKYTASRVDLLFGSNSQLRALAEVYAADD  707

Query  716  AQPKFVQDFVAAWDKVMNLDRFDVR  740
            A+ KFV+DFVAAW KVM+ DRFDV+
Sbjct  708  AKEKFVRDFVAAWTKVMDADRFDVK  732
```
