## Supplementary material for "Genomic analysis of *Mycobacterium brumae* sustains its nonpathogenic and immunogenic phenotype": Data Sheet 3.PDF

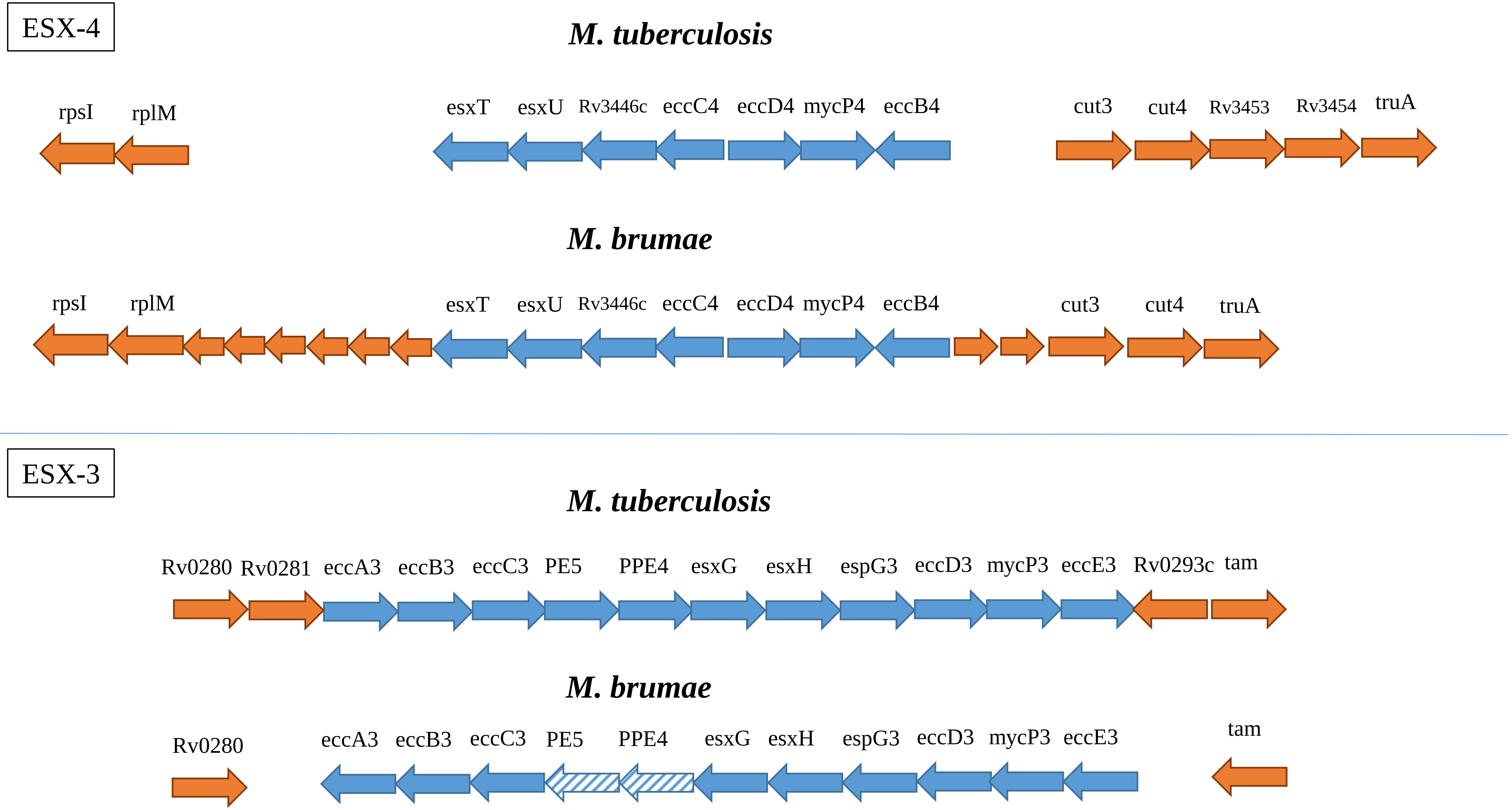


**Supplementary Figure S3.** Comparison of the ESX-3 and ESX-4 clusters in *M. tuberculosis* H37Rv and *M. brumae*. Genes from the cluster are colored in blue and flanquing genes are colored in orange. Stripped arrows indicate the genes found with low similitude (under 70% protein identity).
