## Supplementary material for "Genomic analysis of *Mycobacterium brumae* sustains its nonpathogenic and immunogenic phenotype": Data Sheet 4.PDF

**Supplementary Figure S4. Schematic representation of *M. tuberculosis*, *M. bovis* BCG Connaught and *M. brumae* cell wall structure.** The scheme shows cell wall components present in (A) *M. tuberculosis*, (B) *M. bovis* BCG Connaught and (C) *M. brumae* grown in Middlebrook 7H10 culture medium. Neither the structures nor the whole cell wall is drawn to scale. TMM (trehalose monomycolate), TDM (trehalose dimycolate), PAT (polyacyl-trehalose), DAT (diacyl-trehalose), TAT (triacyl-trehalose), SGL (sulfoglycolipids), PDIM (phthiocerol DIMycocerosates), PGL (phenolic glycolipids), PL (phospholipids), LAM (lipoarabinomannan), LM (lipomannan), PIM (phosphatidyl inositol mannosides).

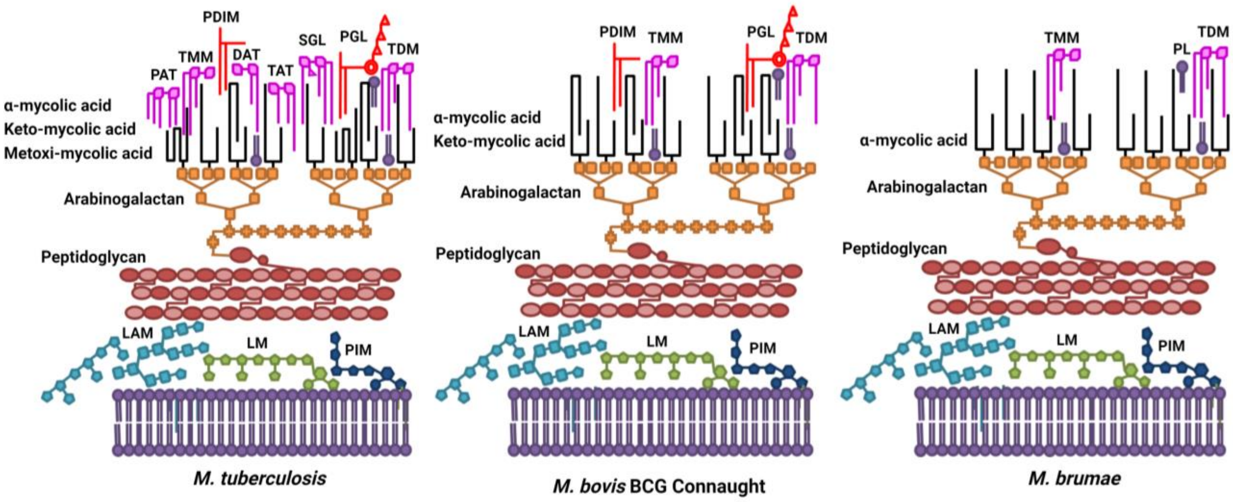
