## Supplementary material for "Genomic analysis of *Mycobacterium brumae* sustains its nonpathogenic and immunogenic phenotype": Data Sheet 5.PDF

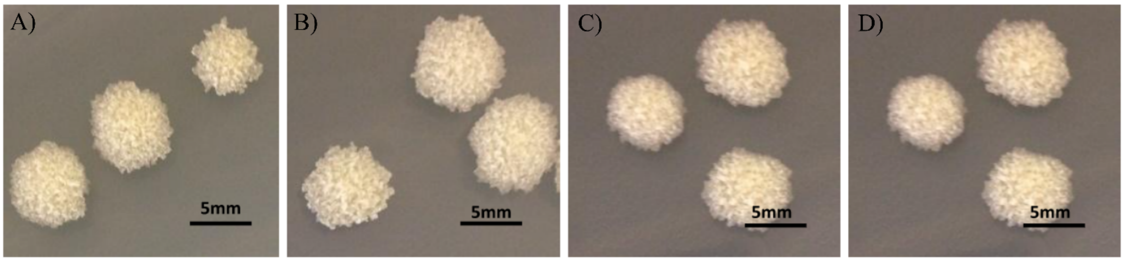


**Supplementary Figure S5.** Colonies of *M. brumae* CR270 (A), CR269 (B), CR142 (C) and CR103 (D) strains grown on Middlebrook 7H10 medium for one week. Pictures were taken with Nikon camera.
