## Supplementary material for "Genomic analysis of *Mycobacterium brumae* sustains its nonpathogenic and immunogenic phenotype": Table 1.DOCX

**Supplementary Table S1. Downloaded genomes used for the phylogenetic analysis.**

| **Species** | **Selected strain** | **Genome size (bp)** | **NCBI Assembly accession number** |
| --- | --- | --- | --- |
| *M. abscessus* | *M. abscessus bolletii* 50594 | 5,270,527 | GCF_000445035.1 |
| *M. avium* | *Mycobacterium avium* subsp. *avium* ATCC 25291 | 4,857,995 | GCF_000174035.1 |
| *M. intracellulare* | *Mycobacterium intracellulare* subsp*. intracellulare* MTCC 9506 | 5,589,007 | GCF_000298095.1 |
| *M. ulcerans* | *Mycobacterium ulcerans* subsp. *shinshuense strain* ATCC 33728 | 5,899,681 | GCF_002355775.1 |
| *M. kansasii* | *M. kansasii* 662 | 6,896,162 | GCF_000523615.1 |
| *M. leprae* | *M. leprae* Br4923 | 3,268,071 | GCF_000026685.1 |
| *M. canettii* | *Mycobacterium canettii* CIPT 140070002 | 4,308,975 | GCF_000343875.1 |
| *M. tuberculosis* | *Mycobacterium tuberculosis* H37Rv | 4,411,532 | GCF_000195955.2 |
| *M. bovis* | *Mycobacterium tuberculosis* variant bovis BCG | 4,410,431 | GCF_001043255.1 |
| *M. confluentis* | *Mycolicibacterium confluentis* JCM 13671 | 5,876,557 | GCF_010729895.1 |
| *M. chitae* | *Mycolicibacterium chitae* NCTC10485 | 5,461,769 | GCF_900637205.1 |
| *M. fallax* | *Mycolicibacterium fallax* | 4,156,821 | GCF_010726955.1 |
| *M. smegmatis* | *M. smegmatis* MC2 155 | 5,521,023 | GCF_000418535.1 |
| *M. fortuitum* | *M. fortuitum fortuitum* DSM 46621 | 6,300,050 | GCF_000295855.1 |
| *M. rutilum* | *M. rutilum* DSM 45405 | 5,987,831 | GCF_900108565.1 |
| *M. phlei* | *M. phlei* RIVM601174 | 5,681,954 | GCF_000257725.1 |
| *M. gilvum* | *M. gilvum* Spyr1 | 5,783,292 | GCF_000184435.1 |
| *M. vanbaalenii* | *M. vanbaalenii* PYR-1 | 6,491,865 | GCF_000015305.1 |
| *M. vaccae* | *M. vaccae* RIVM | 6,223,660 | GCF_000295825.1 |
| *M. insubricum* | *M. insubricum* JCM 16366 | 4,634,331 | GCF_010731615.1 |
| *H. subflava* | *Hoyosella subflava* DQS3-9A1 | 4,863,490 | GCF_000214175.1 |
