## Supplementary material for "Genomic analysis of *Mycobacterium brumae* sustains its nonpathogenic and immunogenic phenotype": Table 6.DOCX

**Supplementary Table S6.** **Susceptibility of the different *M. brumae* strains to different antimicrobials.** A panel of antimicrobials were evaluated against the four strains of *M. brumae* using a Sensititre RAPMYCOI and SLOMYCOI panels. *, susceptibility results were obtained from previously published studies by Luquin *et al*. ([1993](https://doi.org/10.1099/00207713-43-3-405)). The results of the drug susceptibility assays are shown as Minimum Inhibitory Concentration (MIC) (µg/mL). S, sensitive; I, intermediate and R, resistant. n.d., not determined.

| **Drugs** | ***M. brumae* strain** | | | | **Interpretation** |
| --- | --- | --- | --- | --- | --- |
|  | **CR103** | **CR142** | **CR269** | **ATCC 51384** |  |
| Amikacin | 1 | 1 | 1 | 1 | S |
| Amoxicilin/clavulanic acid 2:1 | 2/1 | 2/1 | 2/1 | 2/1 | S |
| Cefepime | 32 | 32 | 32 | 32 | R |
| Cefoxitin | 4 | 4 | 4 | 8 | S |
| Ceftriaxone | 16 | 16 | 16 | 32 | I |
| Ciprofloxacin | 0.12 | 0.12 | 0.12 | 0.12 | S |
| Clarithromycin | 0.06 | 0.06 | 0.06 | 0.06 | S |
| Doxycycline | 0.12 | 0.25 | 0.25 | 0.25 | S |
| Ethambutol | 1-4 | 2-4 | 4 | 2 | S |
| Ethionamide | 2.5 | 2.5 | 2.5 | 2.5 | S |
| Imipenem | 2 | 2 | 2 | 2 | S |
| Isoniazid | 1-2 | 1 | 2-4 | 2 | R |
| Linezolid | 1 | 1 | 1 | 1 | S |
| Minocycline | 1 | 1 | 1 | 1 | S |
| Moxifloxacin | 0.12 | 0.12 | 0.12 | 0.12 | S |
| Rifabutin | 0.5-1 | 0.5-1 | 1 | 0.5 | S |
| Rifampin | 1 | 2 | 2 | 2 | R |
| Streptomycin | 1-2 | 1 | 2 | 1 | S |
| Sulfamethoxazole | 0.12/2.38 | 0.12/2.38 | 0.12/2.38 | 0.12/2.38 | S |
| Tigecyclin | 0.5 | 0.5 | 0.5 | 0.25 | S |
| Tobramycin | 2 | 2 | 2 | 2 | S |
| *p*-Amninosalicylic acid* | n.d. | n.d. | n.d. | n.d. | R |
| Capreomycin* | n.d. | n.d. | n.d. | n.d. | S |
| Cycloserine* | n.d. | n.d. | n.d. | n.d. | S |
| Kanamycin* | n.d. | n.d. | n.d. | n.d. | S |
